# Mucosal innate immune activation as the trigger to *Prevotella* species-induced arthritis in genetically resistant mice

**DOI:** 10.1101/2025.03.18.643707

**Authors:** Yuichi Maeda, Miriam Rabenow, Wei Xiang, Eva Schmid, Lena Amend, Nadine Otterbein, Ilann Sarfati, Ippei Miyagawa, Ilia Gimaev, Leona Ehnes, Till Robin Lesker, Heike Danzer, Kerstin Sarter, Kerstin Dürholz, Fabian Schälter, Lili Bao, Stefan Wirtz, Philipp Arnold, Jörg Hofmann, Daniel Sjöberg, Anna Svärd, Alf Kastbom, Georg Schett, Till Strowig, Mario M. Zaiss

## Abstract

An altered gut microbiota, particularly the expansion of *Prevotellaceae* members, is increasingly implicated in the pathogenesis of rheumatoid arthritis (RA), yet the mechanisms hind this phenomenon remain unclear. Here, we demonstrate that *Palleniella intestinalis*, a member of the *Prevotellaceae* family, induces a 100% arthritis incidence in genetically resistant C57BL/6 mice. Inoculation with *P. intestinalis* modifies gut microbiota ecology, increases intestinal permeability, and selectively activates colonic CD11b⁺CD11c⁺ myeloid cells, facilitating Th17 differentiation and driving joint inflammation. *In vitro*, outer membrane vesicles (OMV) from *P. intestinalis* and *Segatella copri* (formerly known as *Prevotella copri*) prime bone marrow-derived dendritic cells (BMDCs) to drive Th17 differentiation in an IL-6-dependent manner. Similar changes with increase in innate immune cell activation and IL-6 levels were shown in gut biopsies from new-onset RA patients. The transfer of *Prevotellaceae*-derived OMVs or *Prevotellaeae*-primed BMDCs replicates the hightened arthritis incidence in resistant mice, highlighting the critical role of intestinal immune activation in RA.

**Graphical Abstract:** 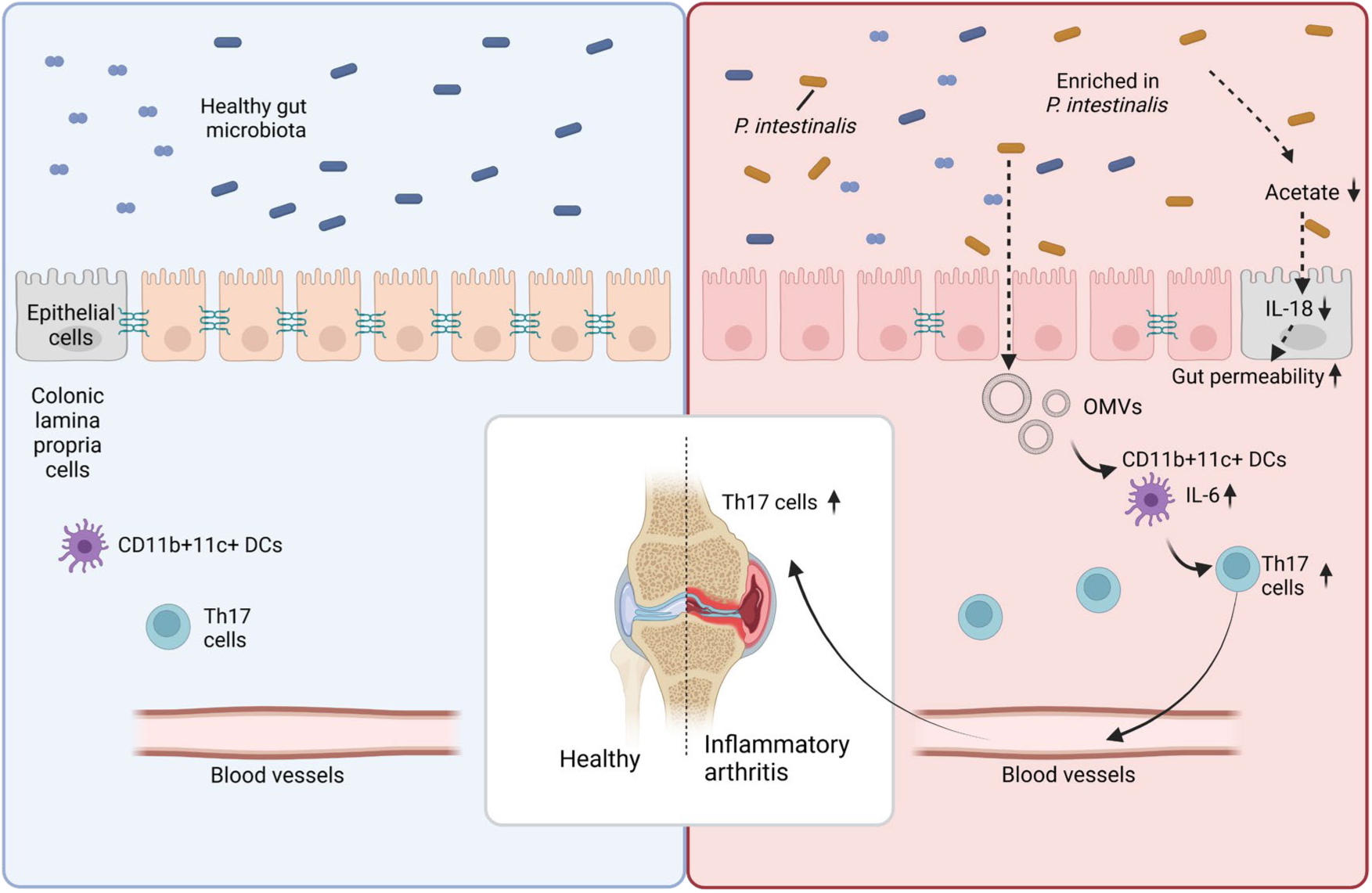

- Colonization with *Paleniella intestinalis* bypasses genetic resistance, consistently inducing arthritis incidence in non-susceptible C57BL/6 mice
- *P. intestinalis* activates colonic CD11b^+^CD11c^+^ myeloid cells, driving systemic Th17 cell differentiation
- In vitro, outer membrane vesicles from *P. intestinalis* or RA-derived *S. copri* RPC01 primed DCs that drive Th17 differentiation in an IL-6-dependent manner
- Transfer of *P. intestinalis*-primed DCs or its outer membrane vesicles alone increases arthritis incidence in non-susceptible C57BL/6 mice

## INTRODUCTION

Rheumatoid arthritis (RA) is a systemic autoimmune disease characterized by synovitis and bone destruction (1, 2). Although significant progress has been made in the treatment of RA, its exact etiology is still not fully understood. While many studies have linked genetic risk alleles to RA, they do not fully explain its incidence (3, 4). A twin study of RA indicated that genetic factors account for only 15–30% of the cases among monozygotic twins (5). This suggests that environmental factors are critical in disease onset, even in genetically predisposed individuals. Among environmental triggers, smoking, hormonal changes, periodontal disease, and intestinal dysbiosis have been proposed as contributing factors to RA development (6–10).

Emerging evidence suggests that the intestinal microbiota is altered in RA patients (11–14). The detection of immunoglobulin A (IgA) anti-citrullinated protein (CCP) antibodies 5–10 years before arthritis onset further supports the hypothesis that RA begins at mucosal sites (15, 16). Additionally, studies have shown that mice or rats raised in germ-free conditions do not develop arthritis unless they are exposed to bacteria (17–19). These findings highlight the importance of the intestinal microbiota in both human and murine arthritis.

Members of the family *Prevotellaceae*, particularly *Segatella copri* (formerly *Prevotella copri*), are enriched in the gut microbiota of patients with early-stage RA (11, 12). Recent studies have shown that the expansion of *Prevotellaceae* in the gut is observed in pre-clinical stages of RA and in individuals genetically predisposed to RA without symptoms (20, 21). We and others have demonstrated that *S. copri* is involved in development of murine models of arthritis (11, 14, 22). However, in non-Westernized countries, *Prevotellaceae* are considered commensal gut microbiota associated with a plant-rich diet due to their ability to degrade plant polysaccharides (23, 24). This raises the question of whether *Prevotellaceae* play a beneficial role or exacerbate autoimmune features in the host. Besides the enrichment of specific bacterial taxa in RA patients, the intestinal epithelial barrier is disrupted in both patients with pre-clinical human arthritis and in collagen-induced murine arthritis (25, 26). However, the mechanisms through which *Prevotellaceae* could contribute to the disruption of the epithelial barrier and activate immune cells in the intestine remain unclear. Furthermore, the specific components of *Prevotellaceae* responsible for immune cell activation during arthritis have not been fully elucidated.

A challenge in studying the effects of human gut-derived *Prevotellaceae* is that these species, e.g., *S. copri*, do not naturally colonize the intestines of specific pathogen-free (SPF) mice. Colonization of *S. copri* in mouse models requires germ-free (GF) conditions or antibiotic (abx) pre-treatment before inoculation (14, 22). Notably, in abx-treated mice, *S. copri* colonization is transient, complicating performing complex disease models. Moreover, GF mice are known to be relatively immunocompromised, even when reconstituted with human gut microbiota (27). Thus, there is a need for a murine model to investigate how *Prevotellaceae* contribute to arthritis under physiological conditions. *Palleniella intestinalis* has been shown to effectively colonize the murine intestine after transfer to SPF mice, efficiently outcompeting other commensal including related *Prevotellaceae* (28). While transfer and successful colonization of *P. intestinalis* has been shown to worsen DSS-induced colitis (29) and bone homeostasis (30), no previous studies have linked this bacterium to arthritis. Since *S. copri* is unable to colonize the gut microbiota of SPF mice without the use of antibiotics, we propose that *P. intestinalis* serves as a valuable model for studying the biological effects of *S. copri* colonization. Notably, *P. intestinalis* and *S. copri* share similar metabolic signatures, such as higher succinate production (29).

In this study, we examined the role of *P. intestinalis* in collagen-induced arthritis (CIA) using C57BL/6 mice, a strain genetically resistant to arthritis (31, 32). We found that inoculation with *P. intestinalis* significantly increased the incidence of arthritis. Our results demonstrated that colonization with *P. intestinalis* shifted the gut microbiota towards a *Prevotellaceae*-dominanted profile, which increased gut permeability and reduced colonic IL-18 levels. This altered environment activated intestinal CD11b^+^CD11c^+^ cells, including dendritic cells (DC), promoting the differentiation of Th17 cells and inducing joint inflammation in an IL-6 dependent manner. Furthermore, outer membrane vesicles (OMVs) derived from *P. intestinalis* were sufficient to induce arthritis in the mice. Interestingly, we also observed innate immune activation and elevated IL-6 levels in the intestines of human individuals with new-onset RA. These findings highlight the crucial role of intestinal innate immune activation in *Prevotellaceae*-induced arthritis in both mice and humans.

## RESULTS

### *P. intestinalis* colonization induces arthritis in mice previously resistant to the disease

We investigated the role of *Prevotellaceae* under physiological conditions and in inflammatory arthritis. For our analysis, we utilized a recently isolated and taxonomically described *P. intestinalis* strain that can colonize SPF mice (29, 33). It has been reported that in the C57BL/6 background, the CIA model exhibits only low incidence rates of approximately 10–40% (31). However, SPF mice colonized with *P. intestinalis* showed incidence rates of up to 100% in three independent experiments (each with n=10), compared to 30-40% in control SPF mice (**Figures 1A and 1B**). Histological analysis of the inflamed paw revealed greater inflammatory cell infiltration in the joints of *P. intestinalis*-inoculated mice (**Figures 1C and 1D**). Additionally, we included SPF mice colonized with *Duncaniella muris*, a member of the *Muribaculaceae* family, which belongs to the same order as *P. intestinalis*. Interestingly, these mice exhibited a low incidence of arthritis, similar to the control group (**Supplemental Figure 1**), suggesting that *P. intestinalis* specifically increased arthritis incidence. We also investigated whether the fecal microbiota composition was altered following the single inoculation of *P. intestinalis.* Fecal samples were collected at three time points: -11 days post collagen type II (CII) immunization (dpi) at the day of inoculation, 1 and 19 dpi. Before colonization (= -11 dpi), gut microbiota α-diversity was similar in both groups (**Figure 1E**). However, α-diversity decreased in *P. intestinalis*-inoculated mice at both 1 and 19 dpi. Notably, β-diversity, analyzed using principal coordinates analysis (PCoA), revealed distinct clustering of *P. intestinalis* and control microbiota communities following inoculation (**Figure 1F**). At the genus level, all *P. intestinalis*-inoculated mice exhibited a dominance of *Prevotellaeae*, while this was absent in control mice (**Supplemental Figure 2**). These results suggest that a single inoculation of *P. intestinalis* significantly altered gut microbiota composition, leading to a *Prevotellaceae*-dominated gut environment.

**Figure 1.**
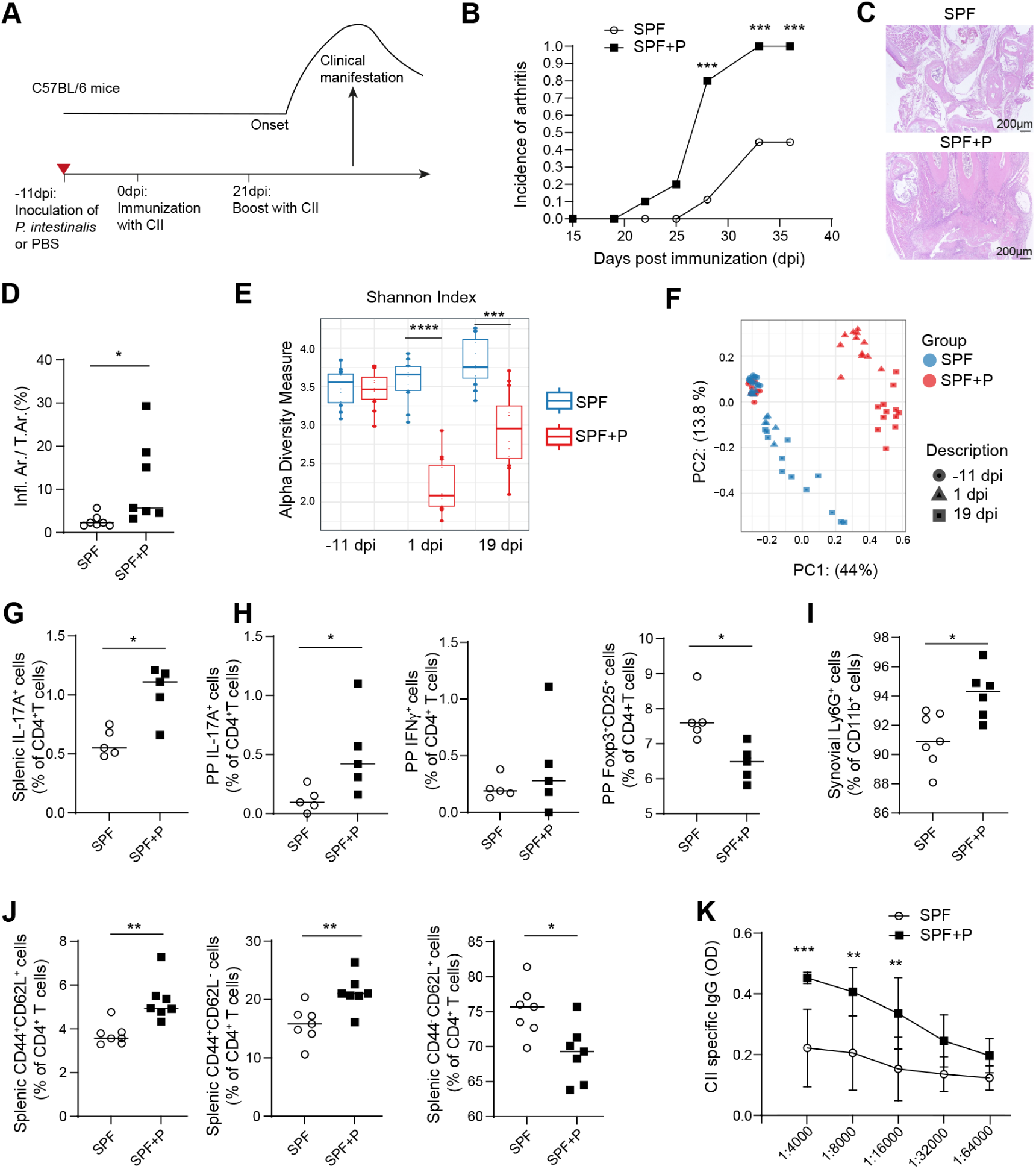
*P. intestinalis* colonization induces arthritis in mice previously resistant to the disease. (A) Experimental outline of collagen-induced arthritis (CIA) in mice inoculated with *P. intestinalis* or control. (B) Incidence of CIA in mice inoculated with *P. intestinalis* (n = 10; SPF+P) versus control (n = 10; SPF). The x-axis represents the days post-first CII immunization (dpi). (C) Histopathological analysis of the front paw at 37 dpi, with one representative image shown (H&E staining). (D) Quantification of the inflamed area as a percentage of the total area in the front paw from H&E staining. (E) α-Diversity analysis of gut microbiota, assessed using the Shannon index. (F) β-Diversity analysis (PCoA) of gut microbiota using Bray-Curtis distances, with multivariate analysis of variance. Fecal samples were collected at three time points: -11, 1, and 19 dpi, corresponding to pre-inoculation, 11 days post-inoculation, and 30 days post-inoculation, respectively. (G) Percentage of IL-17A^+^ cells in CD4^+^ cells from the spleen of *P. intestinalis* (n = 5; SPF+P) versus control (n = 5; SPF) mice at 23 dpi. X axis represents serum dilution factors. (H) Percentage of IL-17A^+^, IFNγ^+^, and Foxp3^+^CD25^+^ cells in CD4^+^ cells from Peyer’s patches (PP). (I) Frequency of Ly6g^+^ cells among CD11b^+^ synovial cells. (J) Proportions of CD44^+^CD62L^+^, CD44^+^CD62L^-^, and CD44^-^CD62L^+^ cells among CD4^+^ cells in the spleen. Immune cells were analyzed by flow cytometry at 37 dpi. (K) ELISA results showing a significant increase in CII-specific IgG serum autoantibody levels at 37 dpi (n = 5 in each group). All data are representative of at least two independent experiments. Statistical significance was determined by two-way ANOVA for (B) and (K), and unpaired t-test for (E, G–J). *p < 0.05, **p < 0.01, ***p < 0.001.

### *P. intestinalis* triggers local and systemic immune responses that precede the onset of arthritis

We analyzed immune cell populations in the spleen, Peyer’s patches, and synovial cells using flow cytometry at two time points: 23 dpi, representing the early stages of arthritis, and 37 dpi, when fully developed arthritis was present in *P. intestinalis*-inoculated mice. At 23 dpi, we observed an increased Th17 cells in the spleen and a trend toward increase in Th17 cells in the Peyer’s patches of *P. intestinalis*-inoculated mice (**Figures 1G and Supplemental Figure 3A**). At 37 dpi, Th17 cells remained elevated in Peyer’s patches, while Th1 cells did not significantly increase (**Figures 1H and Supplemental Figure 3B**). In contrast, the proportion of Foxp3^+^CD25^+^ regulatory T cells among CD4^+^ T cells in the Peyer’s patches was decreased in *P. intestinalis*-inoculated mice (**Figure 1H**). Additionally, there was a tendency for increased Th17 cells in the spleen of these mice (**Supplemental Figure 3C**). Next, we examined immune cell populations in the synovial tissues at both time points. *P. intestinalis*-inoculated mice exhibited an increased proportion of Ly6G⁺ cells among CD11b⁺ cells at both early and late stages (**Figures 1I and Supplemental Figure 3D**). The mean thickness of the front paw was also significantly increased in these mice (**Supplemental Figure 3E**). We further analyzed the proportions of central memory, effector memory, and naive CD4^+^ T cells in the spleen at 37 dpi. Interestingly, *P. intestinalis*-inoculated mice exhibited a significant increase in central memory and effector memory T cells, along with a reduction in naive CD4⁺ T cells (**Figure 1J**). Additionally, serum type II collagen-specific IgG levels were higher at both time points in *P. intestinalis*-inoculated mice (**Figures 1K and Supplemental Figure 3F**). These results suggest that *P. intestinalis* drives a shift toward Th17 cell-mediated immune responses in the early phase of arthritis.

Furthermore, altered gut permeability preceded the onset of arthritis. When sequentially measuring gastrointestinal permeability using FITC-Dextran after immunization, *P. intestinalis*-inoculated mice exhibited significantly higher mean fluorescence intensity (MFI) of FITC in the serum already at the early phase of arthritis (25 dpi), (**Figure 2A**). A similar trend was observed at peak of disease (37 dpi), suggesting a persistent increase in gut permeability. To determine whether increased gut permeability contributed to arthritis onset, we pooled all the mice for analysis and stratified the mice into two groups based on the presence or absence of arthritis at 37 dpi. We then compared their MFI of FITC at 25 and 37 dpi. Mice that developed arthritis by 37 dpi had significantly higher serum FITC-MFI levels at both time points compared to those without arthritis (**Figure 2B**). Additionally, serum anti-*P. intestinalis* specific IgG antibodies were elevated at early time points (**Figure 2C**), further supporting the hypothesis that increased gut permeability in the early phase contributes to arthritis development. Fluorescence *in situ* hybridization (FISH) analysis revealed that intestinal bacteria began invading colonic epithelial cells as early as 25 dpi in *P. intestinalis*-inoculated mice, whereas no such invasion was observed in the control group. By 37 dpi, bacterial invasion was markedly increased in *P. intestinalis*-inoculated mice (**Figure 2D**). Our previous study found that epithelial barrier function in ileum was altered in early stages of CIA mice(25). Therefore, we investigated whether the expression levels of tight junction-related genes in the ileum were also affected in our model. On 23 dpi, *P. intestinalis*-inoculated mice exhibited significantly lower expression of *Claudin-1* and *Claudin-8* (**Figure 2E**) with a trend towards decreased *Occludin* expression. At the later time point, the expression levels of *Zo-1*, *Occludin*, and *Claudin-1* in the ileum were significantly lower in *P. intestinalis*-inoculated mice (**Figure 2F**). Additionally, *Zo-1* expression in the colon was reduced at the later time point (**Figure 2G**). To further investigate whether *P. intestinalis* altered epithelial cell function, we conducted *in vitro* experiments using Caco-2 cells co-cultured with heat-killed *P. intestinalis*, *Bacteroides thetaiotaomicron* (*B. theta*) as a control commensal bacterium, or phosphate-buffered saline (PBS) as a control. After 12 hours of co-culture, we measured the barrier index and found that *P. intestinalis* significantly impaired epithelial barrier integrity in contrast to *B. thetaiotaomicron* (*B. theta*) (**Figure 2H**). These findings suggest that *P. intestinalis* disrupts the epithelial barrier in both the ileum and colon at an early after immunization, leading to increased gut permeability and a higher incidence of arthritis.

**Figure 2.**
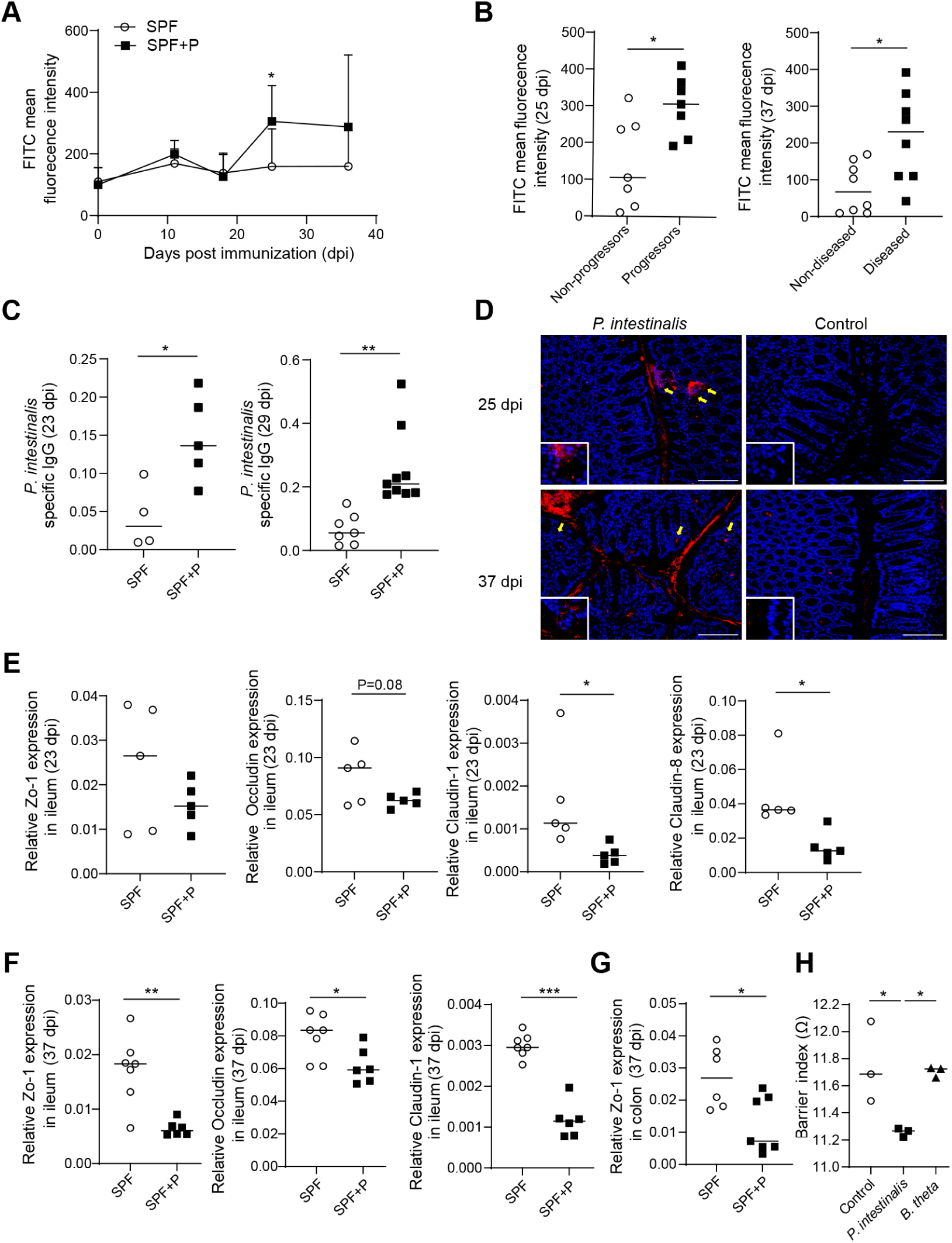
*P. intestinalis* triggers local and systemic immune responses that precede the onset of arthris. (A) Time course of intestinal barrier permeability in CIA mice inoculated with *P. intestinalis* (n = 7; SPF+P) or control (n = 9; SPF). (B) Intestinal barrier permeability at 25 and 37 dpi in mice with (Progressors, Diseased) or without (Non-progressors, Non-diseased) arthritis at 37 dpi. (C) *P. intestinalis*-specific IgG levels in serum at 23 and 29 dpi. (D) Representative FISH imaging of the colon in both groups. Yellow arrows indicate bacterial invasion of colonic epithelial cells at 25 and 37 dpi (n=5). Scale bars, 128µm. (E, F) mRNA expression levels of Zo-1, Occludin, Claudin-1, and Claudin-8 in the ileum at 23 dpi (E), and Zo-1, Occludin, and Claudin-1 at 37 dpi (F). (G) mRNA expression levels of Zo-1 in the colon at 37 dpi. (H) Transepithelial electrical resistance measurement of Caco-2 cell monolayers stimulated with heat-killed *P. intestinalis*, *B. thetaiotaomicron* (B. theta), or control. All data are representative of at least two independent experiments. Statistical differences were determined by two-way ANOVA (A), unpaired t-test (B, C, E–G), and one-way ANOVA (H). *p < 0.05, **p < 0.01, ***p < 0.001.

### CD4⁺ T cells play a pivotal role in *P. intestinalis*-triggered arthritis

To determine whether CD4⁺ T cells play a critical role in *P. intestinalis*-induced arthritis, we induced CIA following the same protocol as in Figure 1A. We then sorted CD4⁺ T cells from the spleen of *P. intestinalis*-treated and control mice and transferred them into CD4 knockout (KO) mice before inducing CIA (**Figure 3A**).

**Figure 3.**
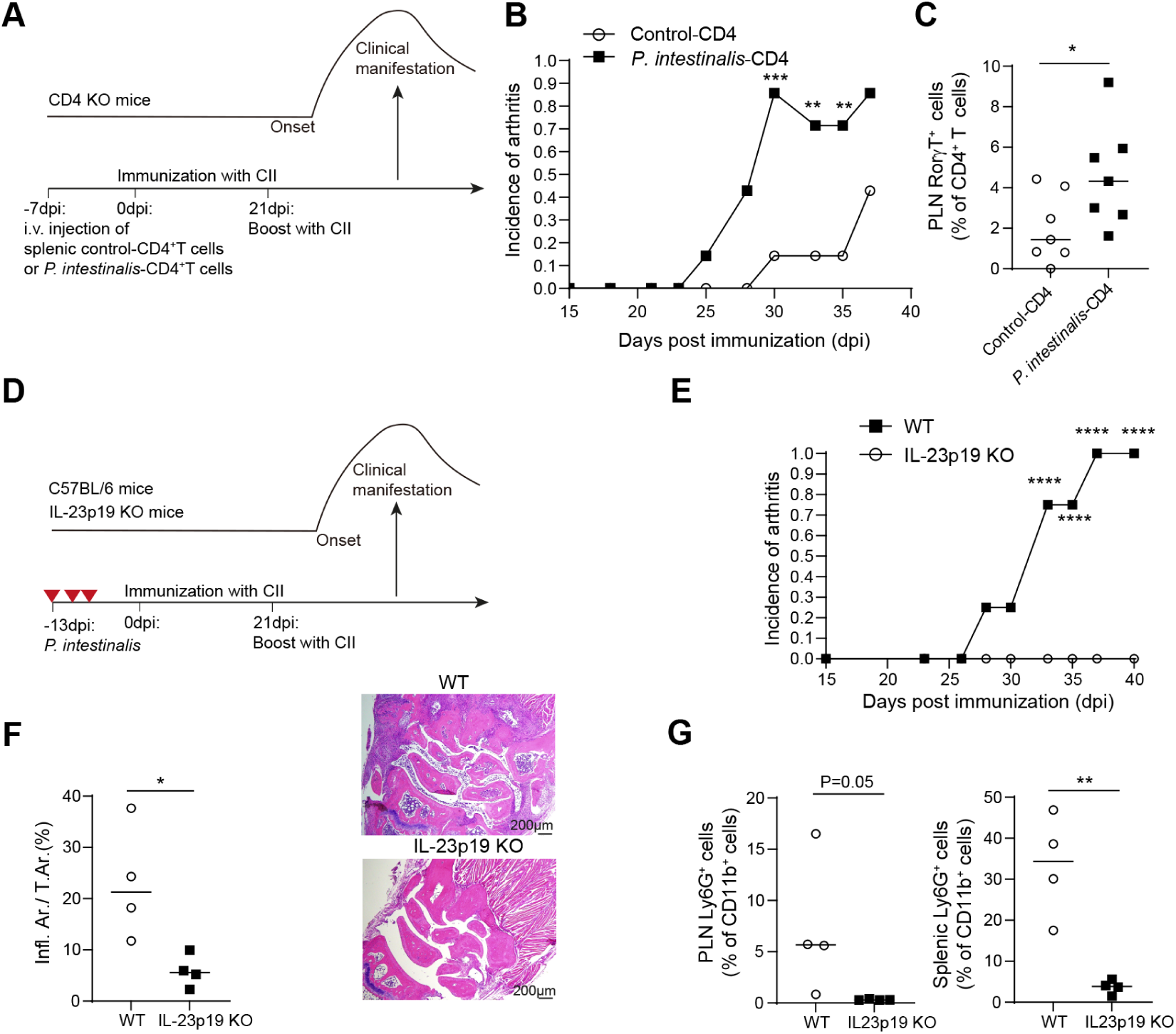
CD4⁺ T cells play a pivotal role in *P. intestinalis*-triggered arthritis. (A) Experimental outline of CIA in CD4 KO mice transferred with splenic CD4+ T cells derived from *P. intestinalis*- or control-inoculated mice. (B) Incidence of CIA in CD4 KO mice transferred with splenic CD4^+^ T cells treated with *P. intestinalis* (n = 7; *P. intestinalis*-CD4) versus control (n = 7; Control-CD4). The x-axis indicates days post-inoculation (dpi). (C) Ratio of RorγT^+^ cells among CD4^+^ T cells in popliteal lymph nodes (PLN). (D) Schematic overview of CIA in WT mice or IL-23p19 KO mice receiving *P. intestinalis*. (E) Incidence of CIA in WT (n = 5) and IL-23p19 KO mice (n = 4) inoculated with *P. intestinalis*. (F) Quantification of the inflamed area as a percentage of the total area in the front paw from H&E staining. Representative images are shown. (G) Ratio of Ly6G^+^ cells among CD11b^+^ cells in PLN and spleen. All data are representative of at least two independent experiments. Statistical differences were determined by two-way ANOVA (B, E) and unpaired t-test (C, F, G). *p < 0.05, **p < 0.01, ***p < 0.001, ****p < 0.0001.

Interestingly, CD4⁺ T cells from *P. intestinalis*-inoculated mice induced a significantly higher incidence of arthritis when transferred into CD4 KO and heterozygous mice (**Figure 3B**). Additionally, the mean paw thickness was greater in these recipient mice compared to those that received CD4⁺ T cells from control mice (**Supplemental Figure 4A**). Immune cell population analysis revealed an increased proportion of RORγT⁺ cells among CD4⁺ T cells in the draining popliteal lymph nodes, but not in the spleen, of mice that had received CD4⁺ T cells from *P. intestinalis*-inoculated donors (**Figures 3C and Supplemental Figure 4B**). Given the observed increase in Th17 cells in *P. intestinalis*-inoculated mice, we next investigated whether Th17 cells are essential for arthritis induction. To test this, we used IL-23p19 KO mice, which lack Th17 cell differentiation and function. Both wild-type (WT) C57BL/6 mice and IL-23p19 KO mice were orally inoculated with *P. intestinalis* and subjected to CIA induction to assess arthritis incidence and severity (**Figure 3D**). We found that WT mice exhibited a higher incidence of arthritis and increased paw thickness, whereas IL-23p19 KO mice did not develop arthritis (**Figure 3E**). Consistent with this, histological analysis showed no inflammatory cell infiltration in the joints of IL-23p19 KO mice (**Figure 3F**). Furthermore, the proportion of Ly6G⁺ neutrophils among CD11b⁺ cells in the popliteal lymph nodes and spleen of IL-23p19 KO mice was significantly reduced (**Figure 3G**).

These findings demonstrate that Th17 cells play a crucial role in the induction of *P. intestinalis*-triggered arthritis.

### *Prevotella*-driven susceptibility to arthritis is associated with IL-6 expression in mice and RA patients

To characterize the differences in colonic inflammation between *P. intestinalis*-inoculated and control mice, we quantified various cytokines in the colon using LegendPlex and ELISA. IL-6 levels were significantly higher in the colon of *P. intestinalis*-inoculated mice, whereas IL-18 levels decreased at 23 dpi (**Figure 4A**). TNF-α levels, however, was unchanged. In contrast, systemic IL-6 and IL-18 levels in the serum did not differ between the two groups, suggesting a locally changed cytokine milieu in the gut (**Figure 4B**). Notably, previous studies have reported that colonic IL-18 levels decrease following *P. intestinalis* inoculation during the early phase of experimental colitis, consistent with our findings (29). A previous study demonstrated that reduced acetate levels modulate IL-18 expression via GPR43 and GPR109A (34). Hence, we hypothesized that changes in short-chain fatty acid (SCFA) levels in the colon might regulate IL-18 expression. To test this, we analyzed SCFA levels in both the colon and serum. Acetate levels were significantly lower in *P. intestinalis*-inoculated mice, while levels of propionic acid, butyric acid, and valeric acid remained unchanged (**Figures 4C**, **4D and Supplemental Figure 5**). These findings suggest that acetate may play a role in modulating IL-18 levels in the colon of *P. intestinalis*-inoculated mice. The role of IL-18 in epithelial cell function is controversial. Some studies suggest that IL-18 promotes gut epithelial integrity, whereas others indicate that it exacerbates colitis (35–37). Additionally, the role of IL-18 in the epithelial barrier during arthritis is unknown. Therefore, we investigated whether recombinant IL-18 (rIL-18) supplementation could reduce arthritis incidence in *P. intestinalis*-inoculated mice. We administered rIL-18 intraperitoneally three times a week from 4 – 25 dpi in both *P. intestinalis*-treated and control mice (**Figure 4E**). Interestingly, rIL-18 significantly reduced arthritis incidence in *P. intestinalis*-inoculated mice (**Figure 4F**). Furthermore, rIL-18 treatment lowered serum CII-specific IgG levels (**Figure 4G**) and improved *Zo-1* expression in the ileum (**Figure 4H**). Histological analysis also showed that rIL-18 ameliorated synovial inflammation (**Figure 4I**). In summary, IL-18 plays a crucial role in maintaining epithelial barrier integrity, and its deficiency - likely caused by reduced acetate levels in the colon - contributes to increased arthritis susceptibility.

**Figure 4.**
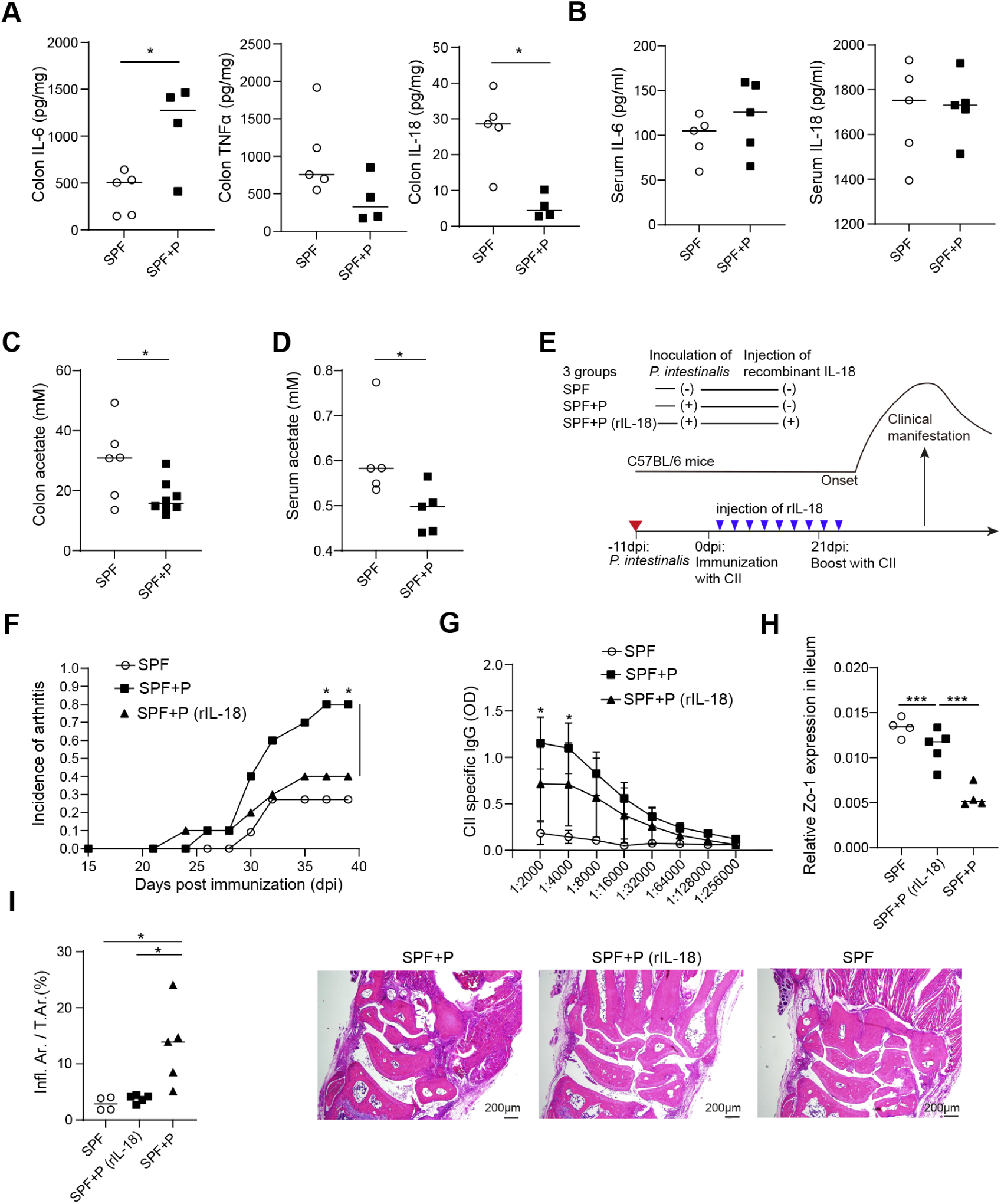
Supplementation of rIL-18 improved arthritis and epithelial barrier. (A) Cytokine levels measured in colon tissue from *P. intestinalis*-treated mice (SPF+P) and control mice (SPF) at 23 dpi. (B) Serum concentrations of IL-6 and IL-18 in mice. (C, D) Acetic acid concentrations in colon contents (C) and serum (D) of *P. intestinalis*-inoculated and control mice. (E) Schematic overview of CIA mice treated with or without rIL-18 in the presence of *P. intestinalis*. (F) Incidence of CIA in mice inoculated with *P. intestinalis* (n = 10, SPF+P) vs. *P. intestinalis* with rIL-18 (n = 10, SPF+P (rIL-18)) vs. control (n = 10, SPF). (G) ELISA results indicate a significant reduction in CII-specific IgG serum autoantibody levels in *P. intestinalis*-inoculated CIA mice treated with rIL-18. (H) mRNA expression levels of *Zo-1* in the ileum at 37 dpi. (I) Histopathological analysis of the front paw at 37 dpi. The proportion of the inflamed area relative to the total area is quantified, along with representative H&E-stained histological images. All data are representative of at least two independent experiments. Statistical significance was determined using an unpaired *t*-test (A–D, I), two-way ANOVA (F, G), or one-way ANOVA (H). *p < 0.05, ***p < 0.001.

To determine whether the observed imbalance of IL-6–Th17 signaling and IL-18 deficiency in the intestine also occurs in human RA, we performed mRNA sequencing on ileal biopsy samples and 16S rRNA amplicon sequencing on fecal samples (**Figure 5A**). The study included 18 RA patients and 9 healthy controls (HC), with the RA group comprising 8 new-onset RA patients and 10 established RA patients (**Supplementary Table 1**).

**Figure 5.**
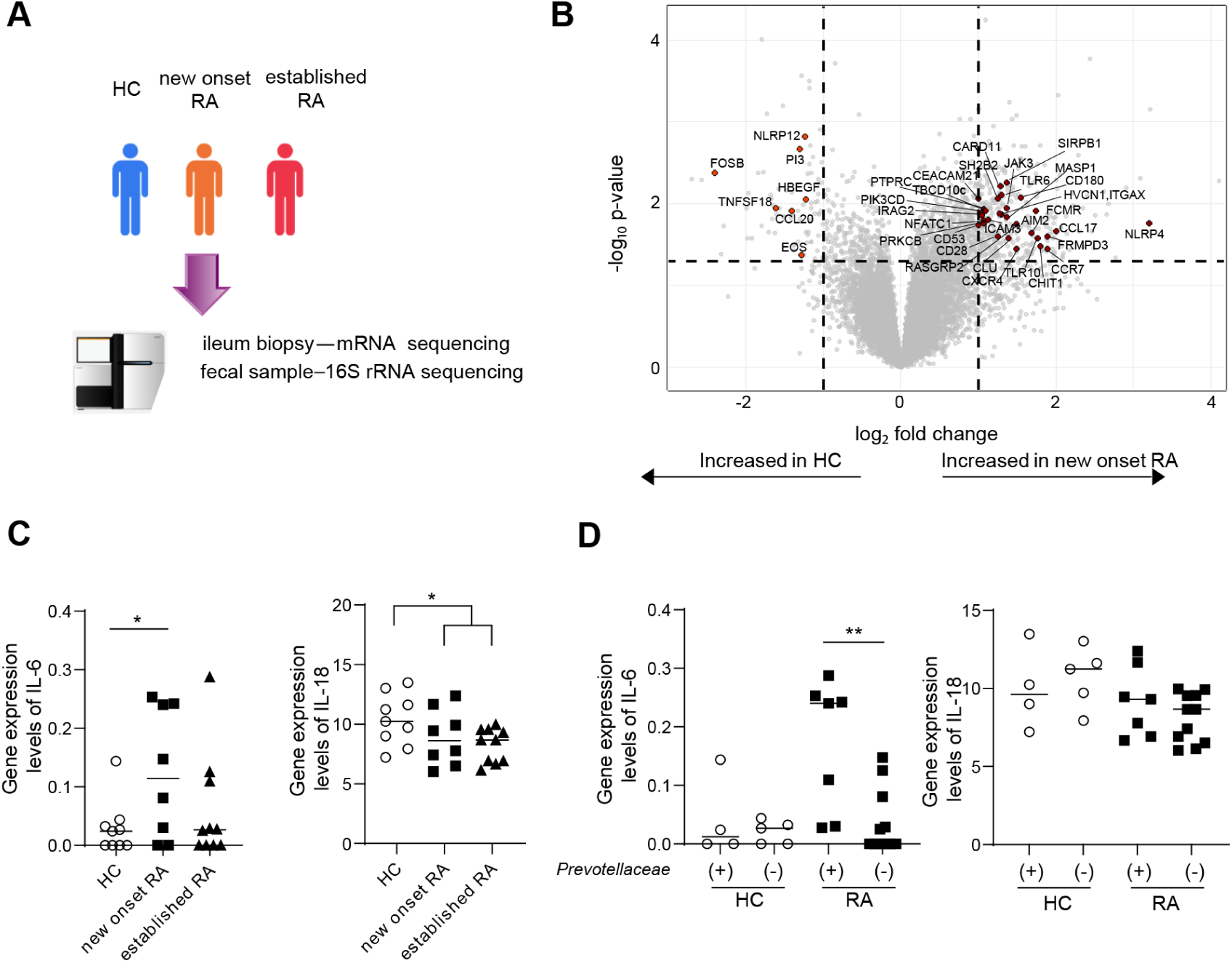
*Prevotella*-driven susceptibility to arthritis is associated with IL-6 expression in mice and RA patients. (A) Graphical summary of mRNA sequencing from ileal biopsies and 16S rRNA sequencing from fecal samples in patients with new-onset RA (n = 8), established RA (n = 10), and healthy controls (n = 9, HC). (B) Volcano plots depicting differentially expressed genes in mRNA sequencing of patients with new-onset RA compared to healthy controls. Genes associated with innate immune activation are highlighted in red. (C) *Gene_fpkm* expression levels of *IL6* and *IL18* in the ileum of patients with new-onset RA, established RA, and HC. (D) *Gene_fpkm* expression levels of *IL6* and *IL18* in RA patients and HC, stratified by the presence (*Prevotellaceae* (+)) or absence (*Prevotellaceae* (-)) of *Prevotellaceae* in the gut. Statistical significance was determined using an unpaired *t*-test (C, D). *p* < 0.05 (*), *p* < 0.01 (**).

In new-onset RA patients, numerous upregulated genes related to innate immune activation and antigen presentation were identified (**Figure 5B**). Additionally, markers of intestinal conventional dendritic cell subset 2 (cDC2) including CD11b, CD11c, and CD1c as well as genes associated with T cell activation, were elevated in new-onset RA patients (**Supplemental Figure 6**). IL-6 expression tended to be higher in RA patients than in healthy controls, being most pronounced in the early stages of RA (**Figure 5C**). In contrast, IL-18 gene expression was significantly lower in RA patients compared to controls (**Figure 5C**). We also examined whether intestinal *Prevotellaceae* influenced IL-6 expression. Notably, RA patients in the presence of *Prevotellaceae* in their feces exhibited higher IL-6 expression in the ileum compared to those without *Prevotellaceae* (**Figure 5D, Supplementary Table 2**). In summary, our findings reveal molecular gut changes in RA resembling increased IL-6 levels and decreased IL-18 levels. This imbalance mirrors the cytokine profile observed in *P. intestinalis*-triggered murine arthritis, suggesting a potential mechanistic link between gut microbiota composition and RA pathogenesis.

### *P. intestinalis*-induced IL-6 expression in gut myeloid cells promotes Th17 differentiation

We hypothesized that activation of innate immune cells in the intestine is essential for Th17 cell induction following epithelial barrier disruption. To investigate this, we examined whether innate immune activation occurred in CIA mice. RNA sequencing of the ileum under SPF conditions after CIA induction revealed significant upregulation of genes associated with innate immune activation, DC activation, and antigen presentation, including antimicrobial peptides such as defensins (**Figure 6A**).

**Figure 6.**
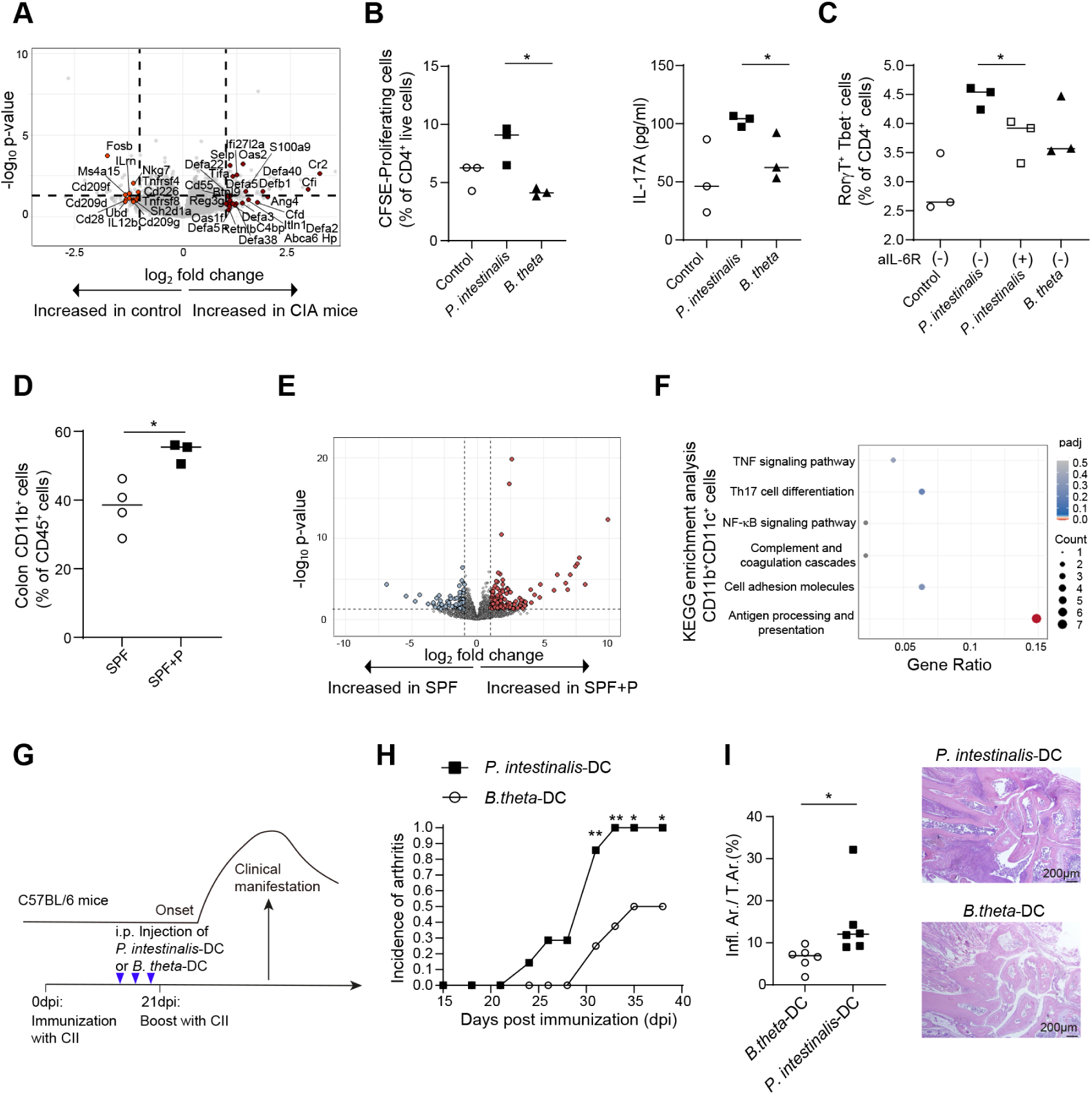
*P. intestinalis*-induced IL-6 expression in gut myeloid cells promotes Th17 differentiation. (A) Volcano plots showing differentially expressed genes in ileal mRNA sequencing of CIA mice. Mice were sacrificed at 5 dpi, and ileal samples were analyzed for mRNA sequencing. Genes associated with innate immune activation are highlighted in red. (B) CFSE proliferation assay of CD4⁺ T cells cocultured with *P. intestinalis*-, *B. thetaiotaomicron* (*B. theta*)-, or control-stimulated dendritic cells (DCs). IL-17A levels in the supernatant were measured by ELISA. (C) Proportion of *RorγT⁺ Tbet⁻* cells among CD4⁺ T cells after coculture of naive T cells with *P. intestinalis* (± anti-IL-6 receptor antibody), *B. theta*, or control-stimulated bone marrow-derived dendritic cells (BMDCs). (D) Ratio of CD11b⁺ cells among CD45⁺ cells in the lamina propria of the colon in *P. intestinalis*-inoculated (SPF+P) and control (SPF) mice. (E) Volcano plot of differentially expressed genes in colonic CD11b⁺CD11c⁺ cells from *P. intestinalis*-inoculated (SPF+P) vs. control (SPF) mice. Upregulated genes in SPF+P are represented by red dots, while downregulated genes are in blue dots. (F) KEGG enrichment scatter plot of CD11b⁺CD11c⁺ cells in *P. intestinalis*-inoculated and control mice. (G) Schematic overview of CIA mice injected with BMDCs stimulated by *P. intestinalis* or *B. theta*. (H) Incidence of CIA in mice injected with BMDCs stimulated by *P. intestinalis* (n = 7, *P. intestinalis*-DC) vs. *B. theta* (n = 7, *B. theta*-DC). The X-axis represents dpi. (I) Histopathological analysis of the front paw at 37 dpi. The inflamed area relative to the total area is quantified, along with representative H&E-stained histological images. All data are representative of at least two independent experiments. Statistical significance was determined using an unpaired *t*-test (B–D) or two-way ANOVA (H). *p* < 0.05 (*), *p* < 0.01 (**).

To assess whether *P. intestinalis*-stimulated DCs promote a Th17 phenotype, we co-cultured CD4⁺ T cells from C57BL/6 mice with *P. intestinalis*-stimulated DCs for 5 days. Upon exposure to *P. intestinalis*-stimulated DCs, T cells exhibited increased proliferation and secreted higher levels of IL-17A compared to controls (**Figure 6B**). We further tested whether IL-6 blockade would reduce Th17 differentiation in the presence of *P. intestinalis*-stimulated DCs. Hence, naive T cells were co-cultured with *P. intestinalis*-stimulated DCs in the presence of an IL-6 receptor antibody. IL-6 blockade significantly reduced the proportion of RorγT⁺ cells among CD4⁺ T cells (**Figure 6C**).

Next, we examined whether intestinal innate immune cells were activated by *P. intestinalis*. To do this, we analyzed the myeloid cell population in the colonic lamina propria shortly after the second immunization, during the early phase of arthritis. Notably, the proportion of CD11b⁺ cells among CD45⁺ cells - including intestinal DCs and macrophages - increased in the intestine (**Figure 6D**). We sorted CD45⁺CD11b⁺CD11c⁺ myeloid cells from the colonic lamina propria, a population known to promote Th17 differentiation and produce high levels of IL-6 (**Supplemental Figure 7A**)(36, 37). Compared to controls, *P. intestinalis*-inoculated mice exhibited altered gene expression patterns in these myeloid cells (**Figures 6E and Supplemental Figure 7B**). Kyoto Encyclopedia of Genes and Genomes (KEGG) pathway enrichment analysis revealed upregulation of genes related to “antigen processing and presentation,” and “Th17 cell differentiation” in *P. intestinalis*-inoculated mice (**Figure 6F**).

CD11b⁻CD11c⁺ cells exhibited lower expression of inflammatory cytokines such as IL-6, and IL-1β compared to CD11b⁺CD11c⁺ myeloid cells (**Supplemental Figure 7C**). Additionally, the gene expression profile of colonic CD11b⁻CD11c⁺ cells did not distinctly differentiate *P. intestinalis*-inoculated mice from controls (**Supplemental Figure 7D**). These results indicate that *P. intestinalis* specifically activates intestinal CD11b⁺CD11c⁺ myeloid cells, leading to the upregulation of genes that promote Th17 differentiation during the initial phase of arthritis.

We then hypothesized that *P. intestinalis*-primed DC activation is a key initiator of arthritis. To test this, we stimulated BMDCs with heat-killed *P. intestinalis* or *B. thetaiotaomicron* (*B. theta*) and injected them intraperitoneally into C57BL/6 mice before inducing CIA. Mice injected with *P. intestinalis*-stimulated DCs developed a significantly higher incidence of arthritis compared to the controls (**Figures 6G and 6H**). Furthermore, histological analysis of the paw showed increased synovial and inflammatory cell infiltration in *P. intestinalis*-stimulated DCs-injected mice (**Figure 6I**). In summary, *P. intestinalis*-stimulated DCs induce Th17 differentiation through IL-6 signaling, contributing to an increased incidence of arthritis.

### *P. intestinalis* and *Segatella copri* RPC01 from a patient with RA, activate BMDCs and induce IL-6

To investigate how *P. intestinalis* activates DCs, we performed *in vitro* experiments using BMDCs from C57BL/6 mice. The DCs were co-cultured with heat-killed *P. intestinalis*, *B. thetaiotaomicron* (*B. theta*), or PBS as a control. We observed that DCs co-cultured with *P. intestinalis* produced significantly higher levels of IL-6 than those exposed to other bacteria (**Figure 7A**).

**Figure 7.**
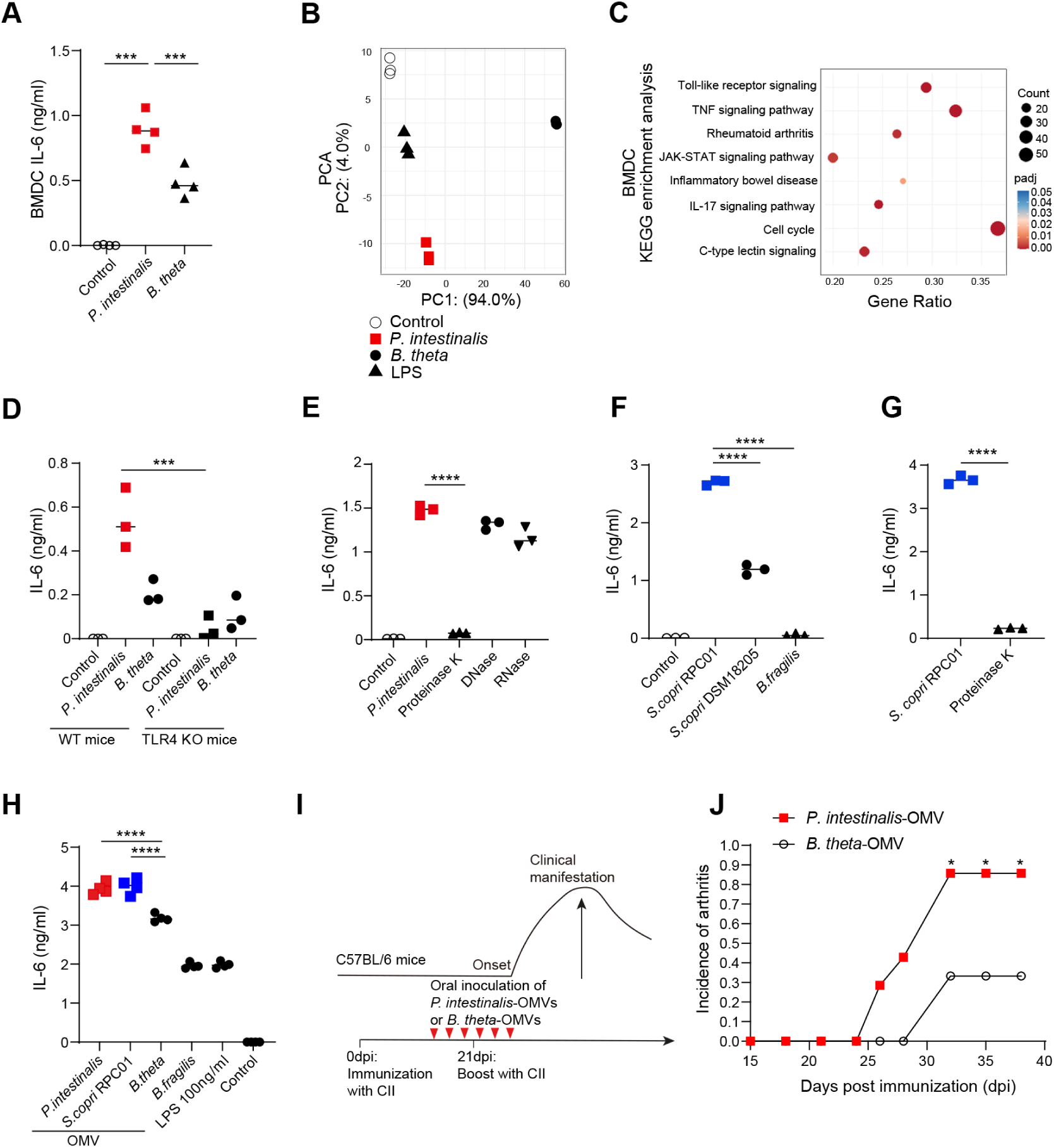
*Prevotellaceae*-derived OMVs induce IL-6 secretion by DCs and transfer CIA susceptibility to resistant mice. (A) IL-6 concentration in the supernatant of BMDCs cocultured with heat-killed *P. intestinalis*, *B. thetaiotaomicron* (*B. theta*), or control, analyzed by ELISA. (B) PCoA of gene expression (FPKM) in BMDCs cocultured with *P. intestinalis*, *B. theta*, LPS, or control. (C) KEGG enrichment scatter plot comparing *P. intestinalis*-stimulated BMDCs with control BMDCs. (D) IL-6 production in the supernatant of BMDCs from WT and TLR4-KO mice cocultured with *P. intestinalis*, *B. theta*, or control. (E) IL-6 levels in BMDCs treated with DNA-, RNA-, or protein-depleted *P. intestinalis* compared to control. (F) IL-6 levels in the supernatant of BMDCs cocultured with *S. copri* RPC01 isolated from an RA patient (*S. copri* RPC01), *S. copri* from a healthy control (*S. copri* DSM 18205), *Bacteroides fragilis* (*B. fragilis*), or control. (G) IL-6 levels in BMDCs treated with *S. copri* RPC01 or protein-depleted *S. copri* RPC01. (H) IL-6 levels in BMDCs treated with outer membrane vesicles (OMVs) derived from *P. intestinalis*, *S. copri* RPC01, *B. theta*, *B. fragilis*, 100 ng/µL LPS, or control. (I) Schematic representation of CIA mice inoculated with *P. intestinalis*-OMV or *B. theta*-OMV. (J) Incidence of CIA in mice inoculated with *P. intestinalis*-OMV (n = 7) vs. *B. theta*-OMV (n = 7). All data are representative of at least two independent experiments. Statistical significance was determined using one-way ANOVA (A, D–F, H), an unpaired *t*-test (G), and two-way ANOVA (J). *p* < 0.05 (*), *p <* 0.01 (**), *p* < 0.0001 (****).

mRNA sequencing of the DCs revealed that the gene expression profile of *P. intestinalis*-stimulated DCs was distinct from that of *B. thetaiotaomicron* (*B. theta*)*-*stimulated, or lipopolysaccharide (LPS)-stimulated DCs, and controls, as shown by PCoA (**Figures 7B and Supplemental Figure 8A**). Specifically, *P. intestinalis*-stimulated DCs exhibited increased expression of IL-6 and IL-1β but not IL-18 compared to *B. thetaiotaomicron* (*B. theta*)-stimulated DCs (**Supplemental Figure 8B**). Importantly, endotoxin levels were similar between *P. intestinalis* and *B. thetaiotaomicron* (*B. theta*), before co-culture (**Supplemental Figure 8C**), indicating that *P. intestinalis* activates DCs and induces IL-6 production independently of LPS stimulation. KEGG pathway enrichment analysis of these DCs revealed upregulation of genes associated with the “IL-17 signaling pathway”, “rheumatoid arthritis”, and “inflammatory bowel disease” in *P. intestinalis*-stimulated DCs (**Figure 7C**). Collectively, these findings suggest that *P. intestinalis*-stimulated DCs strongly promote a Th17 cell response.

Next, we investigated which pattern recognition receptors contribute to IL-6 induction in *P. intestinalis*-stimulated DCs. To this end, DCs from WT and Toll-like receptor 4 knockout (TLR4-KO) mice were co-cultured with *P. intestinalis*, *B. thetaiotaomicron*, or PBS. We found that *P. intestinalis* induced significantly higher TLR4-dependent IL-6 production than *B. thetaiotaomicron* (*B. theta*) (**Figure 7D**). To identify the specific *P. intestinalis* components responsible for IL-6 induction, we treated DCs with *P. intestinalis* pre-treated with DNase, RNase, or proteinase to deplete its DNA, RNA, or protein content, respectively. Notably, protein-depleted *P. intestinalis* failed to induce IL-6 production, suggesting that the cytokine response is protein-dependent (**Figure 7E**).

We then tested whether this observation extended to human-derived *Prevotella* species. Specifically, we used *S. copri* RPC01 isolated from an RA patient, a species belonging to the same former *Prevotellaceae* family as *P. intestinalis*. Remarkably, *S. copri* RPC01 induced significantly higher IL-6 levels in DCs compared to *S. copri* DSM18205 from healthy individuals or compared to stimulation with *Bacteroides fragilis*, used as a control bacterium (**Figure 7F**). The variation in immune activation, such as IL-6 secretion by DCs among different *S. copri* strain, is consistent with previous studies (14, 38). This difference was independent of endotoxin levels (**Supplemental Figure 8D**). Furthermore, *S. copri* RPC01 also triggered IL-6 production in DCs via TLR4 signaling, similar to *P. intestinalis* (**Supplemental Figure 8E**). These findings suggest that *S. copri* and *P. intestinalis* share a conserved protein-dependent mechanism for IL-6 induction in DCs.

### *Prevotellaceae*-derived OMVs induce IL-6 secretion by DCs and transfer CIA susceptibility to resistant mice

To further investigate *P. intestinalis* components responsible for immune activation, we examined OMVs, which are known to modulate host immune responses (39). We hypothesized that *Prevotella*-derived OMVs activate innate immunity and promote IL-6 production. To test this, we isolated OMVs from *P. intestinalis*, *S. copri* RPC01, *B. thetaiotaomicron*, and *B. fragilis* (**Supplemental Figures 9A and 9B**). Notably, OMVs from *P. intestinalis* and RA-derived *S. copri* RPC01 induced significantly higher IL-6 production in DCs compared to those from other commensal bacteria (**Figure 7H**). Finally, we assessed whether *P. intestinalis*-derived OMVs could induce arthritis in mice. Compared to OMVs from *B. thetaiotaomicron* (*B. theta*), *P. intestinalis*-derived OMVs increased both arthritis incidence and paw thickness **(Figures 7I-J and Supplemental Figure 9C**). These results demonstrate that *P. intestinalis*-derived OMVs are sufficient to induce IL-6 production in DCs and contribute to arthritis development in mice.

## DISCUSSION

Increased levels of *Prevotellaceae* in the human intestine of patients with early RA have been reported in several studies (11, 12, 14, 20, 40). However, it remains controversial whether the increased abundance of *Prevotellaceae* in the intestine of early RA patients is a primary phenomenon or a consequence of inflammation (41). Previous reports have shown that patients without genetic susceptibility to RA, such as those lacking HLA-DRB1 risk variants (12) or testing negative for anti-cyclic citrullinated peptide antibody (ACPA), also harbor *Prevotellaceae* in their intestines (40). Therefore, it is crucial to investigate whether *Prevotellaceae*-induced arthritis can occur in animal models that lack genetic susceptibility to arthritis. In the present study, we found that *P. intestinalis* rendered resistant C57BL/6 mice susceptible to arthritis. Mice of the C57BL/6 strain have been reported to exhibit a very low incidence of arthritis (31, 32).

Herein, we also discovered myeloid cell activation in the intestine of patients with new-onset RA. RA patients colonized with *Prevotellaceae* exhibited high IL-6 expression levels in the intestine. IL-6 is a key cytokine involved in the pathogenesis of RA (42). Inhibition of IL-6 is an approved treatment of RA (43, 44), and IL-6 deficient mice have shown reduced immune responses and lower susceptibility to arthritis (45, 46). Notably, in this study, IL-6 receptor blockade reduced RorγT expression in T cells co-cultured with *P. intestinalis*-stimulated DCs. Therefore, IL-6 signaling inhibition may ameliorate RA by interfereing with gut microbiota-induced myeloid cell activation in the gut.

Our results suggest that *P. intestinalis*-induced arthritis follows a three-step process: (1) *P. intestinalis* disrupts the epithelial barrier by reducing intestinal IL-18 levels; (2) it activates innate immune cells, such as CD11b⁺CD11c⁺ cells, in the intestine; and (3) *P. intestinalis*-primed DCs further promote the generation of pathogenic Th17 cells, thereby increasing arthritis incidence.

First, the initial intestinal change involves epithelial barrier disruption due to reduced IL-18 levels, accompanied by decreased acetate concentrations in the colon, mediated by *P. intestinalis*. Although the role of IL-18 in maintaining epithelial integrity is still controversial (35, 47–49), our study demonstrates that supplementation with recombinant IL-18 (rIL-18) reduced arthritis incidence and ameliorated intestinal epithelial barrier dysfunction. Furthermore, RNA sequencing of the ileum from human RA patients revealed decreased IL-18 expression compared to healthy controls. Additionally, acetate levels, which maintain intestinal epithelial barrier dysfunction were reduced in the serum of *Prevotellaceae*-dominant RA patients (unpublished data) and in individuals at risk for RA progression (50). Therefore, increasing acetate and IL-18 levels in RA patients may represent a promising strategy to mitigate gut barrier dysfunction and arthritis. Notably, acetate is the key metabolite of alcohol, which has shown consistent inhibitory effects on arthritis (51).

Second, we demonstrated that the gene expression patterns of CD11b^+^CD11c^+^ cells in the lamina propria of the colon during the early stages of arthritis in *P. intestinalis*-inoculated mice, showing upregulated genes associated with Th17 differentiation and antigen presentation. Furthermore, *P. intestinalis*-stimulated DCs co-cultured with T cells showed increased IL-17A production, proliferative phenotypes, and triggered arthritis incidence in resistant mice. A previous study showed that the depletion of CD11c^+^ cells ameliorated arthritis and reduced osteoclast-associated gene expression (52). These findings highlight that CD11b^+^CD11c^+^ myeloid cells play a central role in triggering *P. intestinalis*-induced arthritis in mice. Notably, the upregulation of innate immune genes, including cDC2 markers, in the intestines of new-onset RA patients further supports DC-driven mechanisms in human RA.

Third, the pathogenic role of Th17 cells in the CIA model has been well established (53, 54) and our study showed that *P. intestinalis* increased Th17 cells in both the gut and spleen during the preclinical and early stages of arthritis. This phenotype aligns with previous studies on *S. copri*-induced arthritis in SKG mice and the CIA model(11, 14). We further confirmed this by adoptively transferring *P. intestinalis*-inoculated CD4⁺ cells into CD4 KO mice, which typically exhibit lower arthritis incidence (55). The recipient mice showed a higher incidence of arthritis and an increased number of Th17 cells in the joints. Additionally, IL-23p19 KO mice that lack the expression of IL-17 and are resistant to arthritis (56, 57), did not develop disease in the presence of *Prevotellaceae*. However, whether Th17 cells from the gut directly migrate to the joints remains unclear. These findings highlight the critical role of Th17 cells in *Prevotellaceae*-induced arthritis in mice.

Translocation of inflammation to the joints may be explained by outer membrane vesicles (OMVs). We enriched OMVs from *P. intestinalis* and *S. copri* RPC01, isolated from the RA patient. These OMVs induced IL-6 production when co-cultured with DCs and contributed to arthritis development. OMVs, produced by both pathogenic and commensal gram-negative bacteria, are known to trigger immune responses and translocate to distant organs (39). Previous research demonstrated that *Fusobacterium nucleatum*-derived OMVs exacerbate arthritis by translocating the FadA protein into the joints (58). However, reports on *Prevotellaceae*-derived OMVs are limited (59–61). This study provides novel insights into the role of *Prevotellaceae*-derived OMVs in arthritis pathogenesis. We also found that *P. intestinalis*-induced DC activation and enhanced IL-6 production were TLR4-dependent, with a protein component mediating this effect. These findings are consistent with the fact that OMVs typically contain LPS and proteins (39). Our findings suggest that TLR4 may recognize *P. intestinalis*-derived OMV LPS or unidentified proteins, leading to increased IL-6 production in DCs. Identifying the effector protein through proteomic analysis combined with mass spectrometry could be a promising next step. Additionally, investigating whether *P. intestinalis*-derived OMVs directly reach the joints may provide key insights into their role in arthritis pathogenesis. In conclusion, our study uncovers a previously unknown mechanism by which *P. intestinalis* disrupts the intestinal epithelial barrier, activates innate immune cells, and drives Th17-mediated arthritis. In RA patients, we observed similar patterns of heightened IL-6-Th17 signaling and reduced IL-18, mirroring our murine model findings. Targeting OMV release from *Prevotellaceae* could open new avenues for preventing arthritis in *Prevotella*-enriched individuals in the future.

## METHODS

### Mice

Wild-type (WT) and CD4 knockout mice used in the study were on the C57BL/6N background. They were all bred and maintained at the animal facilities of the Helmholtz Centre for Infection Research (HZI) under enhanced specific pathogen-free conditions (SPF). They were transported to our animal facility. IL-23p19 knockout mice were purchased from Charles River. TLR4 knockout mice were received from Medizinische Hochschule Hannover. All the experiments were performed with 8-week-old age-matched and gender-matched animals. In *P. intestinalis*-inoculated mice, *Duncaniella muris*-inoculated mice and the control mice were kept in a vinyl isolator separately during the experiment. The mice received water and feed ad libitum. The rooms have a temperature of 22–23°C and a humidity of 50–60%. There is also a 12 h light-dark rhythm in the holding rooms. Animals are kept in type II long cages, with a maximum of five animals. All of the protocols for animal experiments were approved by the local ethics authorities of Regierung von Unterfranken, Germany (55.2.2-2532-2-2073-17).

### Bacterial colonization in mice

*P. intestinalis* or *Duncaniella muris* culture was grown anaerobically (86% N_2_, 10% CO_2_, and 4% H_2_) from frozen glycerol stock in BHI-S (BHI from Oxoid, supplemented with 10% FBS and 0.1% VitK_3_) medium at 37°C for 2 days. All mice were colonized by oral gavage with each bacterium at a dose of 3 × 10^8^ CFU in 200 µl of BHI media. Control mice were orally received 200 ul of PBS.

### Extraction of bacterial DNA & 16S rRNA gene-based murine microbiota analysis

Genomic DNA was extracted using the ZymoBIOMICS 96 MagBead DNA Kit according to the manufacturer’s instructions. 16S rRNA gene amplification of the V4 region (F515/R806) was performed according to an established protocol previously described (Caporaso *et al*. 2011). Briefly, DNA was normalized to 25 ng/µl and used for sequencing PCR with unique 12-base Golary barcodes incorporated via specific primers (obtained from Sigma). PCR was performed using Q5 polymerase (NewEnglandBiolabs) in triplicates for each sample, using PCR conditions of initial denaturation for 30 s at 98°C, followed by 25 cycles (10 s at 98°C, 20 s at 55°C, and 20 s at 72°C). After pooling and normalization to 10 nM, PCR amplicons were sequenced on an Illumina MiSeq platform via 250 bp paired-end sequencing (PE250).

Resulting raw read were demultiplexed by idmp (https://github.com/yhwu/idemp) according to the given barcodes (Caporaso *et al*. 2011). Libraries were processed including merging the paired end reads, filtering the low-quality sequences, dereplication to find unique sequence, singleton removal, denoising, and chimera checking using the USEARCH pipeline version 11.0.667 [[PMID: 26139637]]. In brief, reads merged by fastq_mergepairs command (parameters: maxdiffs 30, pctid 70, minmergelen 200, maxmergelen 400), filtered for low quality with fastq_filter (maxee 1) and singletons using fastx_uniques command (minuniquesize 2). To predict biological sequences (ASV,zOTUs) and filter chimeras we the unoise3 command (minsize 10, unoise_alpha 2), following the amplicon quantification using usearch_global command (strand plus, id 0.97, maxaccepts 10, top_hit_only, maxrejects 250). Taxonomic assignment was conducted by Constax (classifiers: rdp, sintax, blast) using the GreenGenes2 database [PMIDs: 33961008, 37500913] and summarizing into biom-file for the visualization in phyloseq (PMID: 23630581) and downstream analysis.

### Collagen-induced arthritis

CIA was induced in female C57BL/6N mice through subcutaneous injection at the base of the tail with 100 μl containing 0.25 mg chicken type II collagen (Chondrex, Redmond, WA) in complete Freund’s adjuvant (Difco Laboratory, Detroit, MI), containing 5 mg/ml killed *Mycobacterium tuberculosis* (H37Ra). Mice underwent reimmunization after 21 days of intradermal injection in the base of the tail with this emulsion. The paws were evaluated for joint swelling three times per week. Each paw was individually scored using a 4-point scale: 0, normal paw; 1, minimal swelling or redness; 2, redness and swelling involving the entire forepaw; 3, redness and swelling involving the entire limp; 4, joint deformity or ankylosis or both. The thickness of the paw was also analyzed by a caliper.

### Incidence evaluation of arthritis

The incidence of arthritis was calculated by dividing the number of mice with an arthritis score equal to or greater than 1 point by the total number of mice in each experiment. The rate of incidence for each arthritis model was evaluated at each scoring time point.

### In vivo intestinal permeability measurements

For the FITC-Dextran assay, after 4 h fasting of food and water, mice were orally gavaged with 200 μl of FITC-dextran (FD4) (440 mg/kg body weight), and blood samples were collected 4 h later. The concentration of the FITC-dextran was quantified using a fluorimeter with an excitation wavelength of 490 nm and an emission wavelength of 530 nm.

### Histological analysis

Radial or Ulnar bones and inflamed paws with joints were fixed in 4% formalin for 24 h and decalcified in Ethylenediaminetetraacetic acid (EDTA) (Sigma-Aldrich). These samples were embedded in paraffin, sectioned at 4 μm, and stained with H&E. Inflamed areas were quantified using a microscope (Carl Zeiss) equipped with a digital camera and an image analysis system for performing histomorphometry (Osteomeasure; OsteoMetrics)

### Preparation of *P. intestinalis* cultures for ELISA

The strain was maintained inside an anaerobic chamber using BHI-S medium. Bacterial cultures were grown freshly from an overnight inoculation until the exponentially phase and until reaching an optical density (OD) 600 ranging from 0.2 to 0.5. An aliquot of the culture was plated in serial dilutions (10^-4^, 10^-5^, 10^-6^) on BHI from Oxoid, supplemented with 5% sheep blood and 0.1% VitK_3_ and served later on as a quantification control. The rest of the culture was taken out of the chamber, was pelleted by centrifugation and was washed once with PBS. To inactivate the bacterial cells, the pellet was further resuspended in 1% Formalin and kept in the fridge. After 24 h, Formalin was removed by centrifugation, the pellet was washed twice with PBS and the resulting solution was normalized to 4×10^6^ cells/ml by quantifying the colony forming units (CFU) on the previously prepared BHI blood agar plates. Bacterial solutions were kept at -20°C for long-term storage. The identity of all cultures was additionally checked by Sanger sequencing.

### *P. intestinalis* specific IgG levels in the serum

ELISA plates (Microplates, Cat: 3690) were coated with 25 µl of the previously described bacterial solutions per well (resulting in a final concentration of 10^5^ bacterial cells/well) and were incubated overnight at 4°C. The next day, plates were washed twice with PBS solution supplemented with 0.05% Tween 20 (PBST) and were blocked with PBS-T + 5% milk powder for 1h at room temperature (RT) on a shaker. After washing the plates twice with PBST, 25 µl of murine serum samples were added as duplicates to the plates (dilution 1:50 in PBS + 5% milk) and were incubated for 2 h at RT on a shaker. Following washing four times with PBST, 25 µl diluted a goat anti-mouse IgG conjugated to horseradish peroxidase detection antibody (SouthernBiotech cat. no. 61-6520) were added to the plate and were incubated for 1.5 h at RT on a shaker. After washing 3 times with PBST and twice with PBS, TMB mix was added to the wells (25 µl/well) and was incubated for 10 min. After adding 25 µl of 2N H_2_SO_4_, the plates were measured at the ELISA reader at 450 nm against 570 nm as a reference. All samples were measured in duplicates and bacterial controls were carried out on each plate.

### Fluorescence in situ hybridization

The tissues of the large intestine were isolated from C57BL/6N mice at 25 or 37 dpi. For bacterial detection, FISH was performed on formalin-fixed, paraffin-embedded tissue sections using the general bacterial probe EUB338 with the sequence GCT GCC TCC CGT AGG AGT (5′-labelled with Cy3). Briefly, paraffin sections were incubated for 1 hour at 60°C, paraffin was removed with Roti®-Histol (Carl Roth), and sections were rehydrated in a decreasing ethanol series. Slides were then incubated in hybridization-formamide buffer for 20 min at RT, followed by incubation with the EUB338 probe diluted 1:10 in hybridization buffer for 90 min at 46°C in a hybridization oven. Counterstaining was performed with Hoechst 33342 (1:500; Invitrogen), and slides were mounted with fluorescence mounting medium (Dako). The sections were then observed using a DMI 4000 B microscope.

### ELISA for anti-type II collagen antibody

The determination of anti-type II collagen (CII) antibodies was performed as follows. Nunc-Maxisorp ELISA plates (Thermo Fischer Scientific) were coated with chicken CII (Sigma-Aldrich) overnight at 4℃. Following blocking with 1% bovine serum albumin (BSA), sera were applied at the indicated dilution for 2 h. Binding was detected using a goat anti-mouse IgG conjugated to horseradish peroxidase detection antibody (SouthernBiotech), diluted 1:5000 with the appropriate substrate. Data were read using an ELISA reader at 450 nm and 570 nm as a reference wavelength OD with a Magellan Tecan Microplate reader (Tecan Trading AG).

### Tissue isolation

To prepare cells from the spleen, popliteal lymph nodes and Peyeŕs patch, tissue was minced, and mashed through a pre-wetted 70 μm cell strainer. For the spleen, cells were resuspended in 3 ml RBC lysis buffer and incubated for 5 min. The RBC lysis was stopped by adding 10 ml fluorescence activated cell sorting (FACS) buffer, the samples were centrifuged at 1500 rpm for 5 min and the supernatant was discarded. The pellets were used for further experiments.

For the synovial cells, tissues were incubated with RPMI 1640 containing 4% fetal bovine serum (FBS), 2.5 mg/ml collagenase D (Roche), 40 ug/ml of Dnase (Invitrogen) for 30 minutes at 37℃ and 800 rpm on a shaker. Then, take off the supernatant and transfer it in a 50 ml tube through a 70 μm strainer into 10 ml of FACS buffer. In the next step, tissues were incubated with the RPMI containing 4% FBS, 2.5 mg/ml Collagenase/Dispase (Sigma-Aldrich), and 40 μg/ml of Dnase (Sigma-Aldrich) for 30 minutes. Transfer suspension into the same tube through the strainer.

For the cytokine-staining isolated cells were restimulated. Therefore, 10^6^ cells were seeded into a V-bottom 96 well plate in 1× ReMed and incubated for 4 h at 37°C. All samples were washed with PBS, followed by viability staining for 30 min at 4°C in the dark with the Zombie Aqua Fixable Viability Dye. After washing with FACS buffer the cells were incubated with a mix of extracellular binding antibodies and incubated for 30 min in the dark at RT. For intracellular staining, cells were washed with FACS buffer and fixed by using the Foxp3 Fixation and Permeabilization Kit. After fixation the samples were washed with permeabilization buffer and then incubated with 50 μl of the intracellular antibodies, diluted in permeabilization buffer, for 1 h in the dark. The samples were washed with permeabilization buffer and then reconstituted in FACS buffer until sample acquisition. Samples were acquired with a Cytek Northern Lights.

### Tissue isolation (Intestinal lamina propria)

Intestinal lamina propria lymphocytes and myeloid cells were prepared as described previously (PMID: 18716618). In brief, the colon was opened longitudinally, washed to remove fecal content, and incubated in HBSS with 5 mM EDTA at 37°C for 15 min. After washing with PBS, tissues were cut into small pieces and incubated into RPMI 1640 medium containing 4% FBS, 1 mg/ml collagenase D (Roche), 0.1 mg/ml dispase (Sigma-Aldrich) and 40 μg/ml DNase I (Invitrogen) for 25 min at 37°C in a shaking water bath. The digested tissues were resuspended in 5 ml of 30% Percoll (VWR) and then overlaid on 3 ml of 70% Percoll. Percoll gradient separation was performed by centrifugation at 2,000 rpm for 20 min at RT. The lamina propria cells were collected at the interface of the Percoll gradient and washed with PBS containing 2% FBS.

### Flow cytometry

For flow cytometric analyses, cells were stained with antibodies to the following markers in Supplementary Table 3 for further details of antibody clone, dilution factor, provider, target species and application. Organs were harvested, and single-cell suspensions were prepared. For spectral flow cytometry, single stains were prepared on freshly isolated cells of the same tissue. Unless otherwise mentioned, data was acquired on a Cytek Aurora Northern Lights (Cytek) and analyzed using FlowJo software (version 10.8.1).

### Isolation of CD45^+^CD11c^+^ cells from colonic lamina propria

CD45^+^ CD11b^+^ CD11c^+^ or CD45^+^ CD11b^-^ CD11c^+^ myeloid cells from lamina propria of the colon were isolated using **MoFlo Astrios EQ 1 at** 22 and 24 dpi. PE anti-mouse/human CD11b (clone M1/70, biolegend) APC anti-mouse CD11c (clone N418, ThermoFisher) and FITC anti-mouse CD45 (clone 30-F11, Biolegend), anti-mouse CD16/32 (FcγRIII/II, clone 93, biolegend), Zombie Aqua (Biolegend) were used for cell staining. The isolated cells were put in 350 μl of RLTlysis buffer. The total RNA of the cells was isolated using RNeasy Plus Micro Kit (Qiagen). Extracted RNA was transferred to Novogene for RNA sequencing.

### RT-PCR in mice

Tissues were stored in RNA later (Ambion). RNA was extracted according to the manufacturer’s instructions. Gene expression results are expressed as arbitrary units relative to the expression of the housekeeping gene GAPDH. Primer sequences are as follows: GAPDH: 5′-GGG TGT GAA CCA CGA GAA AT-3′ and 5′-CCT TCC ACA ATG CCA AAG TT-3′; ZO-1: 5′-CCA CCT CTG TCC AGC TCT TC-3′ and 5′-CAC CGG AGT GAT GGT TTT CT-3′; occludin: 5′-CCT CCA ATG GCA AAG TGA AT-3′ and 5′- CTC CCC ACC TGT CGT GTA GT-3′; claudin-1: 5′- TCTACGAGGGACTGTGGATG-3′ and 5′- TCAGATTCAGCAAGGAGTCG-3′; claudin-8: 5′- GCCGGAATCATCTTCTTCAT-3′ and 5′-CATCCACCAGTGGGTTGTAG-3′, Zo-1: 5′-; CCACCTCTGTCCAGCTCTTC-3′ and 5′-CACCGGAGTGATGGTTTTCT-3′.

### Cytokine production in the colon

The colonic tissues were cut into 1cm and washed with antibiotics-included PBS and the tissue was incubated with 4% FBS-included RPMI for 12 h. Then, the supernatant was collected and centrifuged for 10 min. The supernatant was used for IL-6 and TNF-α production by Legendplex^TM^ MU Th Cytokine Panel (12-plex; Biolegend) and IL-18 production by ELISA according to the manufactureŕs instructions. Data was adjusted by the weight of the colonic sections.

### Maestro Z impedance measurements

Before cell plating, 100 µL of DMEM containing 2% FBS was added to each well of CytoView-Z 96-well electrode plates (Axion Biosystems, Atlanta, GA, USA). Plates were then docked into the Maestro Z instrument to measure the baseline impedance. Caco-2 cells were subsequently added to the well, and left at RT for 1 h to allow even distribution. The plates with cells were incubated in the Maestro Z for up to 96 h at 37°C / 5% CO_2_ to allow monolayer formation, monitored by resistance measurement. Resistance was measured at 41.5 kHz and 1 kHz, which reflects cell coverage over the electrode and strength of the barrier formed by the cell monolayer, respectively. Measurements were continuously recorded under identical conditions using the Maestro Z instrument. Experimental wells contained either media alone, 7.5 × 10^5^ heat-killed *P. intestinalis*, or 7.5 × 10^5^ heat-killed *B. thetaiotaomicron*, with 25,000 of Caco-2 cells per well. Raw resistance values were normalized to vehicle control and to the value 10 min prior to the treatment. Data analyses at 12 h after the bacterial stimulation were performed in Axis Z software (Axion Biosystems, Atlanta, GA, USA).

### Stimulation of bone marrow-derived DCs

Bone marrow cells were isolated from the femur and tibia of C57BL/6N mice and differentiated in the presence of recombinant murine GM-CSF (Peprotech) for 7 days to generate BMDCs. 3 × 10^6^ of heat-killed *P. intestinalis* or *B. thetaiotaomicron*, *S. copri* RPC01 from an RA patient or *S. copri* DSM 18205, or *B. fragilis* or PBS were prepared. Endotoxin levels were assessed using Pierce™ chromogenes Endotoxin Quant Kit on 1:10 dilutions of the prepared bacterial suspensions. BMDCs (5 × 10^5^) were cocultured with these bacteria for 12 h and the cytokine concentrations of IL-6 in the supernatants were analyzed by ELISA (ThermoFisher). The total RNA of the stimulated BMDCs was extracted using RNeasy Plus Micro Kit (Qiagen). The BMDCs cocultured with heat killed-*P. intestinalis* or *B. thetaiotaomicron,* 10 ng/ml of LPS, and PBS were used for mRNA sequencing. The extracted RNA samples were transferred to Novogene and analyzed mRNA sequencing.

In some experiments, *P. intestinalis* pretreated with 400 μg/ml of DNA-nase, 400 μg/ml of RNA-nase, and 400 μg/ml of protein-nase to deplete the DNA, RNA, and protein, and *S. copri* RPC01 was pretreated with 400 μg/ml of protein-nase was used for cocultured with BMDCs. In other experiments, the indicated volume of heat-killed *P. intestinalis, B. thetaiotaomicron*, or *S. copri* RPC01 or PBS were cocultured with 1.5 x 10^5^ BMDC cells from C57BL/6N mice or TLR4 KO mice. The supernatant of BMDCs were used after 12 h for analyzing IL-6 production by ELISA.

### RNA sequencing

#### RNA quantification

The quantity and quality of the RNA samples were assessed using the following methods: 1% agarose gel electrophoresis to test RNA degradation and potential contamination. Sample purity and preliminary quantitation were measured using Bioanalyzer 2100 and it was also used to check the RNA integrity and final quantitation. *Library construction and quality control of the library for colonic CD11c^+^ cells.* Total RNA was reverse transcribed into the first-strand cDNA. Then the double-strand cDNA was synthesized by LD-PCR amplification (SMART-Seq V4 Ultra Low Input RNA kit for Sequencing 480 Rxns (Cat No. 634893)), followed by the purification with AMPure XP beads and quantification with Qubit. The cDNA samples were then fragmented, end-repaired, A-tailed, and ligated with adaptors (NEB Next® Ultra™ RNA Library Prep Kit). After size selection and PCR enrichment, the RNA library was ready for sequencing. The library was checked with Qubit and real-time PCR for quantification and bioanalyzer for size distribution detection. Quantified libraries will be pooled and sequenced on Illumina platforms, according to effective library concentration and data amount.

#### Library construction and quality control of the library for murine BMDC and human ileum biopsies

Messenger RNA was purified from total RNA using poly-T oligo-attached magnetic beads. First-strand cDNA was synthesized with random hexamer primers, followed by second-strand synthesis. Libraries were constructed using the Novogene NGS RNA Library Prep Set based on the NEB Next® Ultra™ RNA Library Prep Kit, undergoing end repair, A-tailing, adapter ligation, size selection, amplification, and purification.

#### Sequencing

The qualified libraries were sequenced Next Generation Sequencing (NGS) based on Illumina’s Sequencing Technology by Synthesis (SBS)-detection by fluorescence of the nucleotide added during the synthesis of the complementary chain - and in a parallelized and massive way. For colonic CD11c^+^ cells, the Novaseq X Plus sequencing platform was used to sequence the libraries. For murine BMDC or human ileum samples, the Novaseq6000 sequencing system was used to sequence the libraries.The strategy is paired-end 150bp (PE150), 6G raw data per sample. After quality control, reads containing adaptors and low-quality reads were filtered out. Processed reads were mapped to the reference genome with HISAT2. For murine data, GRCm39 was used as reference genome. For human data, GRCh38 was used as reference genome. Raw counts were counted using featureCounts. FPKM was calculated based on the gene length and mapped read counts.

#### Differential expression analysis

Differential expression analysis was performed on raw counts of protein coding genes using the DESeq2 (version 1.42.1) package in R (4.3.3). Sample clustering was performed on rlog transformed data. In Volcano plots, genes with |log2FoldChange| ≥ 1 and a p-value ≤ 0.05 were indicated as up- or downregulated. For mouse experiments (Figure 6F), KEGG enrichment analysis was performed on genes with adjusted p-value ≤ 0.2 and |log2FoldChange| > 0 using the enrichKEGG function of the clusterProfiler (4.10.1) package. For cell culture experiments (Figure 7C), KEGG enrichment analysis was performed on the list of all genes ranked by |log2FoldChange| using the gseKEGG clusterProfiler function with p-value adjustment by Benjamini-Hochberg correction. Ggplot2 (3.5.1) was used for graphical illustrations.

### Activation of T cells by BMDCs

BMDCs (5 × 10^5^) prepared from C57BL/6N mice were incubated with heat-killed *P. intestinalis*, *B. thetaiotaomicron*, or PBS for 12 h. Splenic CD4^+^ T cells from 8-week-old C57BL/6N mice were sorted by magnetic-activated cell sorting (Biolegend). 1 × 10^6^ CD4^+^ T cells from C57BL/6N mice were co-cultured with 5 × 10^5^ bacteria-stimulated DCs for 5 days. The culture supernatant was collected for ELISA to evaluate cytokine production. In the analysis of T cell proliferation, CD4^+^ T cells were labeled with CFSE (15min, 37°C) and stimulated with indicated DCs for 5 days. CFSE negative proliferating cells were analyzed by CytoflexS Cytometer (Beckman Coulter).

In some experiments, splenic naive CD4^+^ T cells were isolated from C57BL/6N mice by Mojosort Mouse CD4 Naive T cell isolation Kit. Then, 1.2 × 10^6^ of the naive CD4^+^ T cells were co-cultured with 3 × 10^5^ bacteria-stimulated BMDCs for 5 days in the presence or absence of 6 μg of anti-IL-6 receptor antibody (AF1830, R&D systems). Following culture, cells were harvested and stained for markers using Zombie NIR (Biolegend) and APC anti-mouse CD4 (clone GK1.5, Biolegend). After fixation and permeabilization, intracellular staining was performed with PE anti-mouse RorγT (clone Q31-378, BD Biosciences), and BV605 anti-human/mouse T-bet (clone 4B10, Biolegend). Flow cytometry data were acquired on a CytoflexS Cytometer.

### Adoptive transfer of CD4^+^ T cells into CD4 knockout mice

CD4^+^ T cells were isolated from the spleen of the C57BL/6N mice that had been inoculated with *P. intestinalis* or control, and immunized with CIA as shown in Figure 3A. Isolation was performed using Mojosort^TM^ mouse CD4 T cell isolation kit, according to the manufacturer’s presentation. The purity of the CD4^+^ T cells was >95%. CD4^+^ T cells (1 × 10^6^) were intravenously transferred to CD4 knockout (CD4^-/-^) or heterozygous (CD4^+/-^) mice at 7 days before first immunization. The mice were reared in the SPF facility in the vinyl isolator. The CD4 KO mice were induced CIA as previously described and were analyzed for incidence of arthritis and immune cell populations.

### IL23p19 knockout mice inoculated with *P. intestinalis*

WT C57BL/6N mice and IL-23p19 KO were cohoused for two weeks. These mice were separately reared into two groups. Both groups were orally inoculated with *P. intestinalis* three separate times, administered at days -13, -10, and -7 relative to days post CII immunization (dpi) as shown in Figure 3D. CIA was performed as previously described and were analyzed for incidence of arthritis and immune cell populations.

### rIL-18 treatment

WT C57BL/6N mice inoculated with *P. intestinalis* were injected with or without rIL-18 (500 ng/mouse) intraperitoneally 3 times a week from 4 to 25 dpi as shown in Figure 4E. The mice of the control group were inoculated with PBS before immunization and injected with PBS. All the mice were induced CIA as described previously and were analyzed for incidence of arthritis and immune cell populations.

### Injection of bacteria-stimulated BMDCs into CIA mice

BMDCs (5 × 10^5^) prepared from C57BL/6N mice were incubated with 3 × 10^6^ of heat-killed *P. intestinalis*, *B. thetaiotaomicron*, for 12 h. These cells were resuspended with 200 μl of PBS and these were intraperitoneally injected into C57BL/6N mice at 14, 19, and 21 dpi and induced CIA.

### Measurement of SCFAs

Four to five replicates of frozen colonic contents (100 mg) or 50 µl of serum were weighed into a 2 ml polypropylene tube. The tubes were kept in a cool rack throughout the extraction. 33% HCl (50 µl for colonic contents or 5 µl for serum) was added and samples were vortexed for 1 min. One milliliter of diethyl ether was added, vortexed for 1 min, and centrifuged for 3 min at 4 °C. The organic phase was transferred into a 2 ml gas chromatography (GC) vial. For the calibration curve, 100 μl of SCFA calibration standards (Sigma) were dissolved in water to concentrations of 0, 0.5, 1, 5, and 10 mM and then subjected to the same extraction procedure as the samples. For GC mass spectrometric (GCMS) analysis, 1 μl of the sample (4–5 replicates) was injected with a split ratio of 20:1 on a Famewax, 30 m × 0.25 mm iD, 0.25 μm df capillary column (Restek). The GC-MS system consisted of GCMS QP2010Plus gas chromatograph/mass spectrometer coupled with an AOC20S autosampler and an AOC20i auto injector (Shimadzu). The injection temperature was 240°C with the interface set at 230°C and the ion source at 200°C. Helium was used as carrier gas with a constant flow rate of 1 ml/min. The column temperature program started at 40°C and was ramped to 150°C at a rate of 7°C/min and then to 230°C at a rate of 9°C/min and finally held at 230°C for 9 min. The total run time was 40 min. SCFA were identified based on the retention time of standard compounds and with the assistance of the NIST 08 mass spectral library. Full-scan mass spectra were recorded in the 25-150 m/z range (0.5 s/scan). Quantification was done by integration of the extracted ion chromatogram peaks for the following ion species: m/z 45 for acetate eluted at 7.8 min, m/z 74 for propionate eluted at 9.6 min, and m/z 60 for butyrate eluted at 11.5 min. GCMS solution software was used for data processing.

### Generating outer membrane vesicles

OMVs were extracted from *P. intestinalis*, *S. copri* RPC01, *B. thetaiotaomicron*, and *B. fragilis* following the procedures. These bacteria were grown in 50 ml of BHI-S medium at 37°C for 2 days under anaerobic conditions, and the supernatant was collected by centrifugation at 4000 g for 30 min at 4°C. This supernatant was further filtered through a 0.22 μm filter to remove larger particles and bacteria. The filtrate was mixed with polyethylene glycol (PEG) 8000 (Roth) solution 1:5 and 3.75 M NaCl solution 1:50 for precipitation vesicles during an overnight incubation at 4°C. Samples were centrifuged at 4000 g for 30 min at 4°C and pellets were resuspended with 1ml of sterile PBS for an ultracentrifugation step (Beckman Optima MAX-YP Ultracentrifuge) at 110,000 g for 2 h at 4°C. After removing the supernatant, OMVs were resuspended in 500 μl of PBS.

### Nanoparticle tracking analysis (NTA)

OMV samples were diluted 100-, 1000- and 10000-fold in sterile Dulbecco’s PBS (Thermo Fisher). 1000 µL of the diluted sample was injected into a ZetaView Twin (Particle Metrix, Germany) measurement chamber. The particles were counted and measured by the device throughout the chamber, and the NTA software determined the concentration of the samples. The dilution that yielded the particle count per frame closest to the device’s target, and thus provided the most reliable measurement, was selected for determining the particle size and concentration of the sample.

### Transmission electron microscopy (TEM)

Negative Stain images were taken as described previously (PMID: 32456348). In brief, 3µl samples were added to a freshly glow discharged continuous carbon grid (Science Service Munich) and then washed with 5 µl of a 2% uranyl acetat solution twice. After air drying, samples were transferred to a JEOL1400 TEM (JEOL, Garching, Germany) and imaged at a nominal magnification of 30.000.

### Inoculation of OMVs into mice and induce CIA

1 x 10^8^ particle numbers determined by NTA of *P. intestinalis*-derived OMVs or *B. thetaiotaomicron*-OMVs were inoculated into female C57BL/6N mice orally at 13, 15, 19, 21, 24, and 27 dpi. The mice were induced CIA as described above. *In vitro* experiment, 5 x 10^8^ particles of OMVs derived from *P. intestinalis*, *S. copri* from RPC01, *B. thetaiotaomicron*, and *B. fragilis* were cocultured with 5 x 10^5^ BMDCs. 12 h later, IL-6 production was analyzed in the supernatant by ELISA.

### Patients and healthy controls

Eighteen patients with RA and 10 healthy controls from the County of Dalarna, Sweden, were included in the “Intestinal involvement in Rheumatiod Arthritis” (IntestRA) study between 2016 and 2019. Patients with known RA were recruited from the rheumatology department’s waiting list, 10 patients with established RA and 8 patients with new-onset RA (disease duration <1 year) were included. All patients had a clinical diagnosis of RA according to a specialist in rheumatology and were positive for IgG anti-CCP in serum. The clinical characteristics of the groups are shown in Supplementary Table 1. As healthy controls, persons participating in a screening study for colorectal cancer were recruited. None of these persons were diagnosed with colorectal cancer during the procedure. First, fecal samples were collected from all study participants shortly before they began the laxative preparation for the colonoscopy. Then, ileum biopsies were collected from all study participants.

### Ileum biopsy of human samples

We performed an ileocolonoscopy intubating at least 5 cm of the terminal ileum, followed by two biopsies with standard biopsy forceps from the ileal mucosa. The samples were immediately placed in a test tube filled with RNA stabilizing solution (RNAlater™ from Thermo Fisher Scientific). They were frozen within one hour after collection and stored at -80°C until further analyses. The samples were transported to our laboratory, and total RNA was extracted using RNeasy Plus Micro Kit (Qiagen).

### 16S rRNA amplicon sequencing of human fecal samples

Ten ng of stool genomic DNA was used in polymerase chain reaction (PCR) amplification of genomic 16S ribosomal RNA V4 regions using the prokaryotic primer pair (515F forward primer: 5′-GTGYCAGCMGCCGCGGTAA−3′; 806R reverse primer: 5′- GGACTACNVGGGTWTCTAAT−3′) containing barcodes on the forward primer 515F (https://earthmicrobiome.org/protocols-and-standards/16s/). The NEBNext Q5 Hot Start Hifi PCR Master Mix (New England Biolabs, Frankfurt am Main, Germany) was used in a reaction employing 25 PCR cycles. The resulting PCR products were purified with AMPure XP Beads (Beckmann Coulter), pooled in equimolar ratios and analyzed by 2 × 151 paired-end sequencing on an Illumina MiSeq device (Illumina). Raw fastq files were imported and analyzed in QIIME2 v2024.5 with DADA2 as the method for quality control, dereplication, and amplicon sequence variant (ASV) table generation. The SILVA database release 138 was used at a 99% similarity cutoff for taxonomic classification. For further analysis, ASV and taxonomic tables were imported into R (version 4.4) as a phyloseq object. Ggplot2 was used for generation of graphical illustrations. Relative abundance of *Prevotellaceae* greater than 0.02% was defined as the presence of *Prevotellaceae*.

### Statistical analysis

Data are expressed as mean ± s.d. unless otherwise indicated in the figure legends. Analysis was performed using unpaired *t*-test for single comparison, or analysis of variance (ANOVA) test for multiple comparisons (one-way or two-way ANOVA followed by Tukey’s or Bonferroni’s multiple comparisons test, respectively). All experiments were conducted at least two times. *P* values of 0.05 were considered significant and are shown as \**p* < 0.05, \*\**p* < 0.01, \*\*\**p* < 0.001, or \*\*\*\**p* < 0.0001. Graph generation and statistical analyses were performed using the Prism version 8 software (GraphPad).

## Data availability

All relevant data are available from the authors upon reasonable request.

## Supplementary Materials

Supplementary Figures 1-9.

Supplementary Table 1-3.

## Acknowledgments

We thank all members of our laboratories at the Medical Clinic 3, Erlangen, Germany, for their support and helpful discussion. Additional thanks goes to the Flow Cytometry & Fluoresence activated Cell Sorting Core team of Uwe Appelt and Markus Mroz, the Optical Imaging Competence Centre (OICE) and their team with Dr. Philip Tripal for imaging and the complete team of the the Preclinical Experimental Animal Centre (PETZ) at the Franz Penzoldt Centre (FPZ) as the central animal facility of the Faculty of Medicine at FAU and the University Hospital Erlangen. We thank Prof. Dr. Gregor Fuhrmann for NTA measurement. TLR4 KO mice were kindly provided by Prof. Dr. Ulrich A. Maus. We thank Novogene for performing the mRNA sequencing. This study was supported by grants from the Deutsche Forschungsgemeinschaft (DFG, German Research Foundation) via projects DFG499424281, DFG-RU2886-Project A01 as well as DFG-CRC1181-Project-No. B07. This study was further funded by the Interdisciplinary Center for Clinical Research, Erlangen (IZKF) (project number P155) at the Universitätsklinikum Erlangen, Germany.

## Author contributions

Y.M. performed the experiments and analyzed the data. M.M.Z., T.S. and Y.M. planned and directed the overall project and wrote the manuscript. M.M.Z., T.S. and G.S. supervised the project. M.R., W.X., E.S., I.S., L.A, I.M., I.G., L.E., H.D., K.D., F.S., L.B., P.A., and J.H. performed the experiments. N.O., T.R.L., and S.W. analyzed the data. A.S., D.S., and A.K. planned and performed the human ileum biopsies. K.S. has provided support for approval of animal license. All authors approved of the final manuscript.

## Material availability

All material generated in this paper will be made available by the lead contact upon request.

