## Supplemental Figures for "Mucosal innate immune activation as the trigger to *Prevotella* species-induced arthritis in genetically resistant mice"

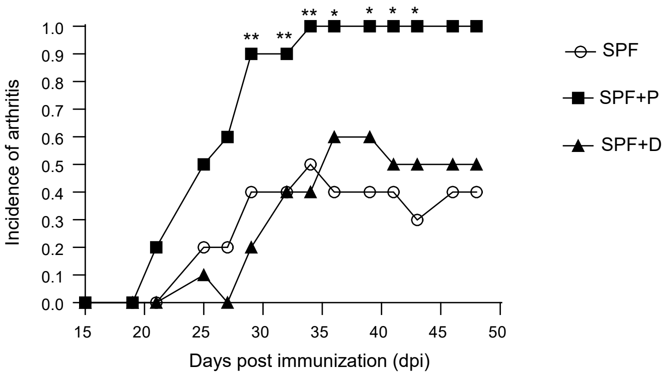


**Figure S1:** ***P. intestinalis*** **specifically increases  arthritis incidence.**
Incidence of CIA in mice inoculated with *P. intestinalis* (n = 10; SPF+P) vs. *Duncaniella muris* (n = 10; SPF+D) vs. control (n = 10; SPF). The X-axis represents dpi. Statistical significance was determined using two-way ANOVA. $p < 0.05$ (*), $p < 0.01$ (**)

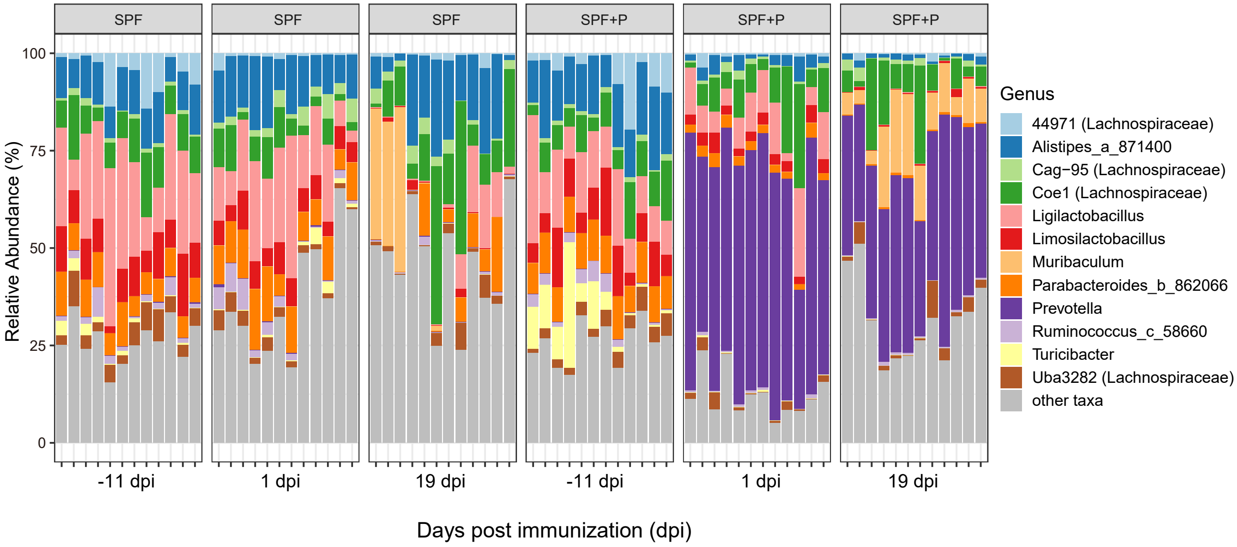


**Figure S2: Colonization of *P. intestinalis* in SPF mice.**
Bacterial composition of fecal samples from *P. intestinalis*-inoculated (SPF+P) and control (SPF) mice. Samples were collected at -11, 1, and 19 dpi. Gut microbiota composition is shown at the genus level.

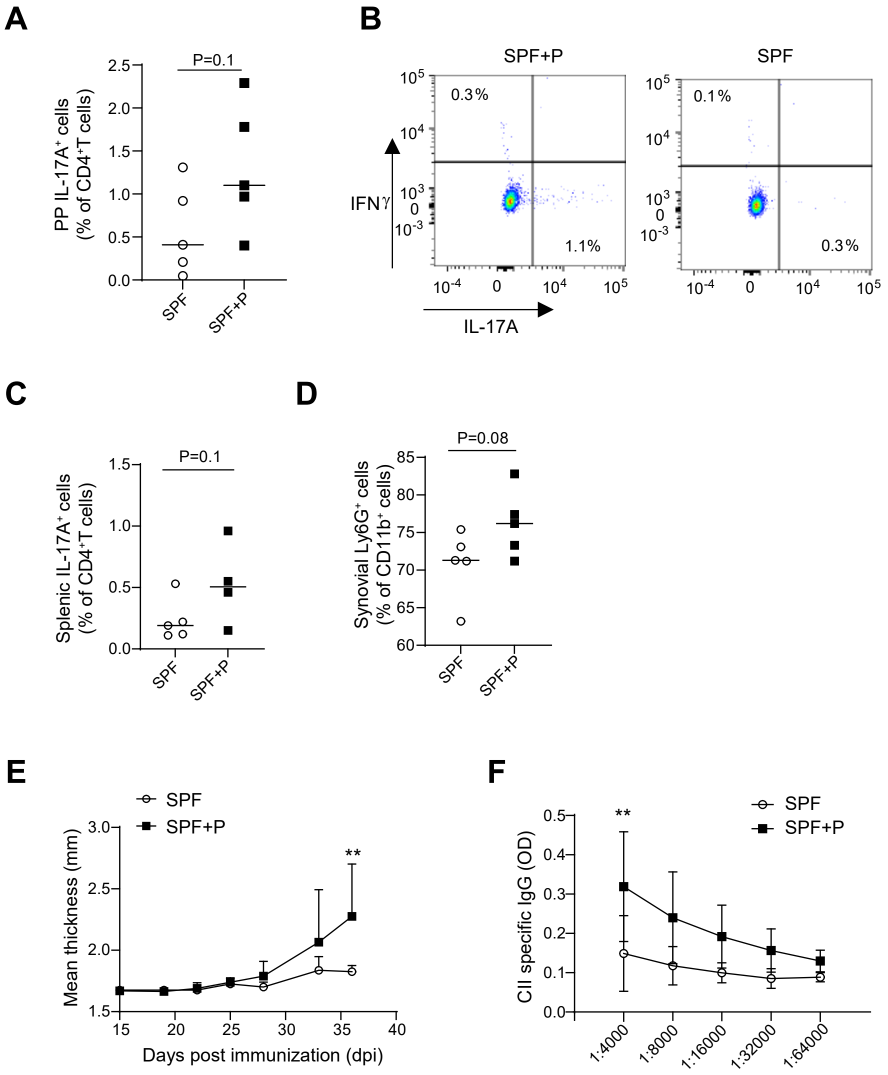


**Figure S3: *P. intestinalis* increases Th17 cells.**
(A) Ratio of IL-17A$^+$ cells among CD4$^+$ T cells in Peyer's patches (PP) from *P. intestinalis*-inoculated (n = 5, SPF+P) and control (n = 5, SPF) mice at 23 dpi. (B) Representative flow cytometry gating strategy for identifying IL-17A$^+$ and IFNγ$^+$ cells among CD4$^+$ T cells in the PP. (C) Ratio of IL-17A$^+$ cells among CD4$^+$ T cells in the spleen at 37 dpi. (D) Ratio of Ly6G$^+$ cells among CD11b$^+$ cells in synovial tissue at 23 dpi. (E) Mean paw thickness in CIA mice inoculated with *P. intestinalis* vs. control. The X-axis represents dpi. (F) ELISA results showing a significant increase in CII-specific IgG serum autoantibody levels in *P. intestinalis*-inoculated CIA mice at 23 dpi (n = 5). Statistical significance was determined using an unpaired *t*-test (A, C, D) or two-way ANOVA (E, F). $p < 0.01$ (**).

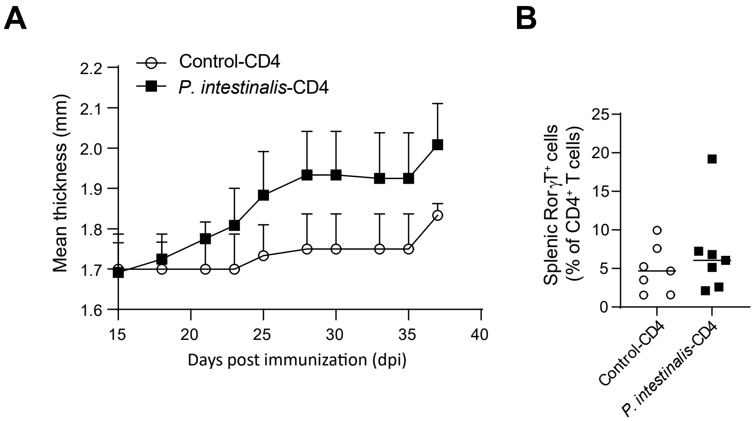


**Figure S4: CD4⁺ T cells play a pivotal role in *P. intestinalis*-triggered arthritis**

(A) Mean paw thickness in CIA mice (CD4 KO) following the transfer of splenic $CD4^+$ T cells treated with *P. intestinalis* (*P. intestinalis*-CD4) vs. control (Control-CD4). (B) Ratio of *RorγT*$^+$ cells among $CD4^+$ T cells in the spleen. Statistical significance was determined using two-way ANOVA (A) and an unpaired *t*-test (B).

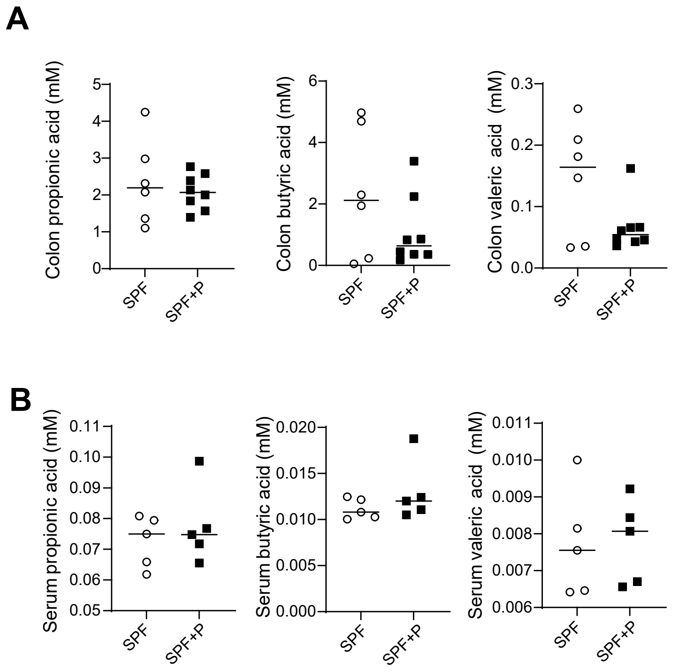


**Figure S5: SCFA production in *P. intestinalis*-inoculated mice.**
(A) Concentration of propionic, butyric, and valeric acid in colon contents of *P. intestinalis*-inoculated (SPF+P) and control (SPF) mice. (B) Concentration of propionic, butyric, and valeric acid in serum of *P. intestinalis*-inoculated (SPF+P) and control (SPF) mice. Statistical significance was determined using an unpaired *t*-test.

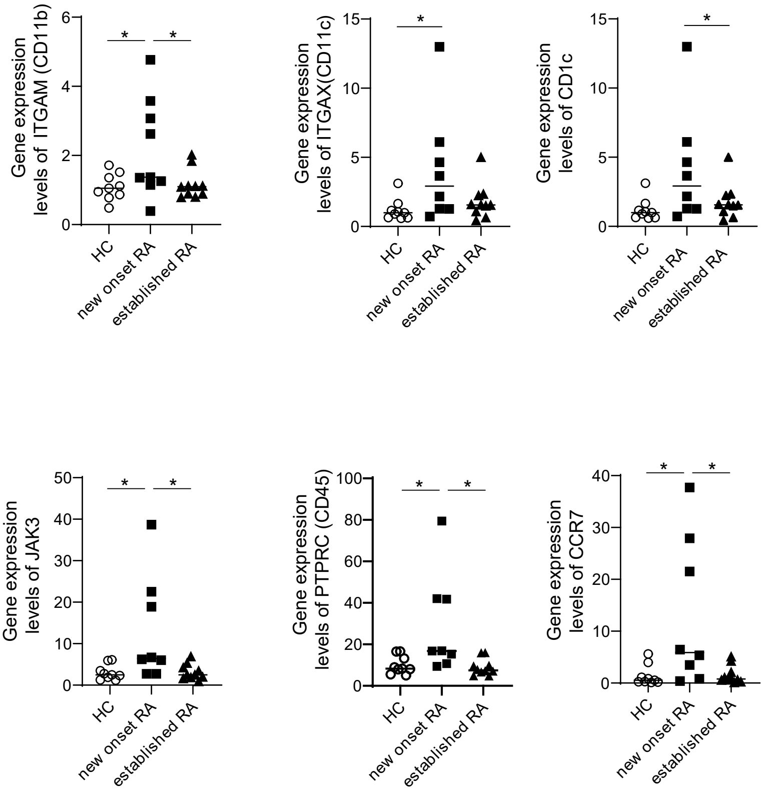


**Figure S6: Gene expression levels in human ileum biopsies.**
Gene_FPKM levels of *ITGAM* (CD11b), *ITGAX* (CD11c), *CD1C*, *JAK3*, *PTPRC* (CD45), and *CCR7* from ileal mRNA sequencing in patients with new-onset RA, established RA, and healthy controls (HC). Statistical significance was determined using one-way ANOVA. $p < 0.05$ (*).

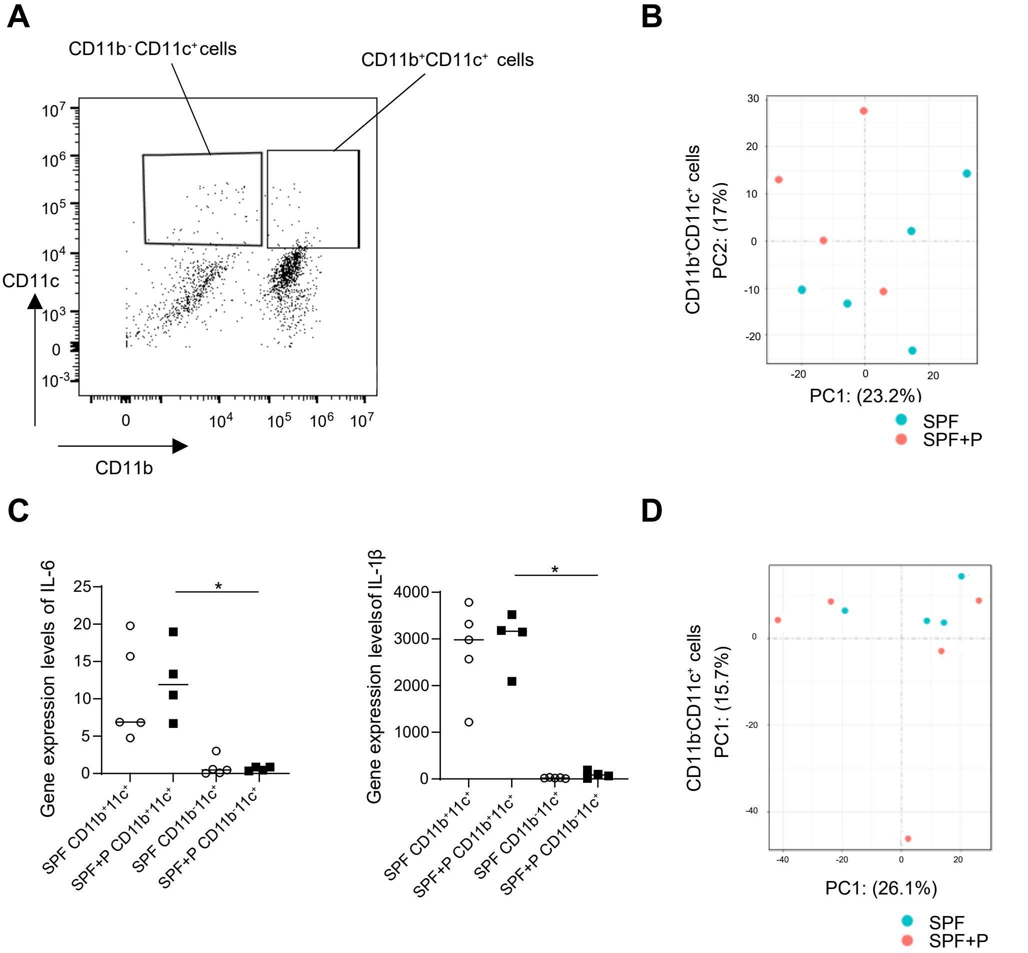


**Figure S7: Gene expression levels in CD11c$^+$ cells in lamina propria.**

(A) Gating strategy for identifying CD11b$^+$CD11c$^+$ and CD11b$^-$CD11c$^+$ cells among CD45$^+$ live cells in the colon. (B) PCoA of gene expression (FPKM) in CD11b$^+$CD11c$^+$ cells from the lamina propria of the colon. (C) Gene expression levels of *IL-6* and *IL-1β* in CD11b$^+$CD11c$^+$ and CD11b$^-$CD11c$^+$ cells from *P. intestinalis*-inoculated (SPF+P) and control (SPF) mice. (D) PCoA of gene expression (FPKM) in CD11b$^-$CD11c$^+$ cells from the colonic lamina propria of *P. intestinalis*-inoculated (SPF+P) and control (SPF) mice.

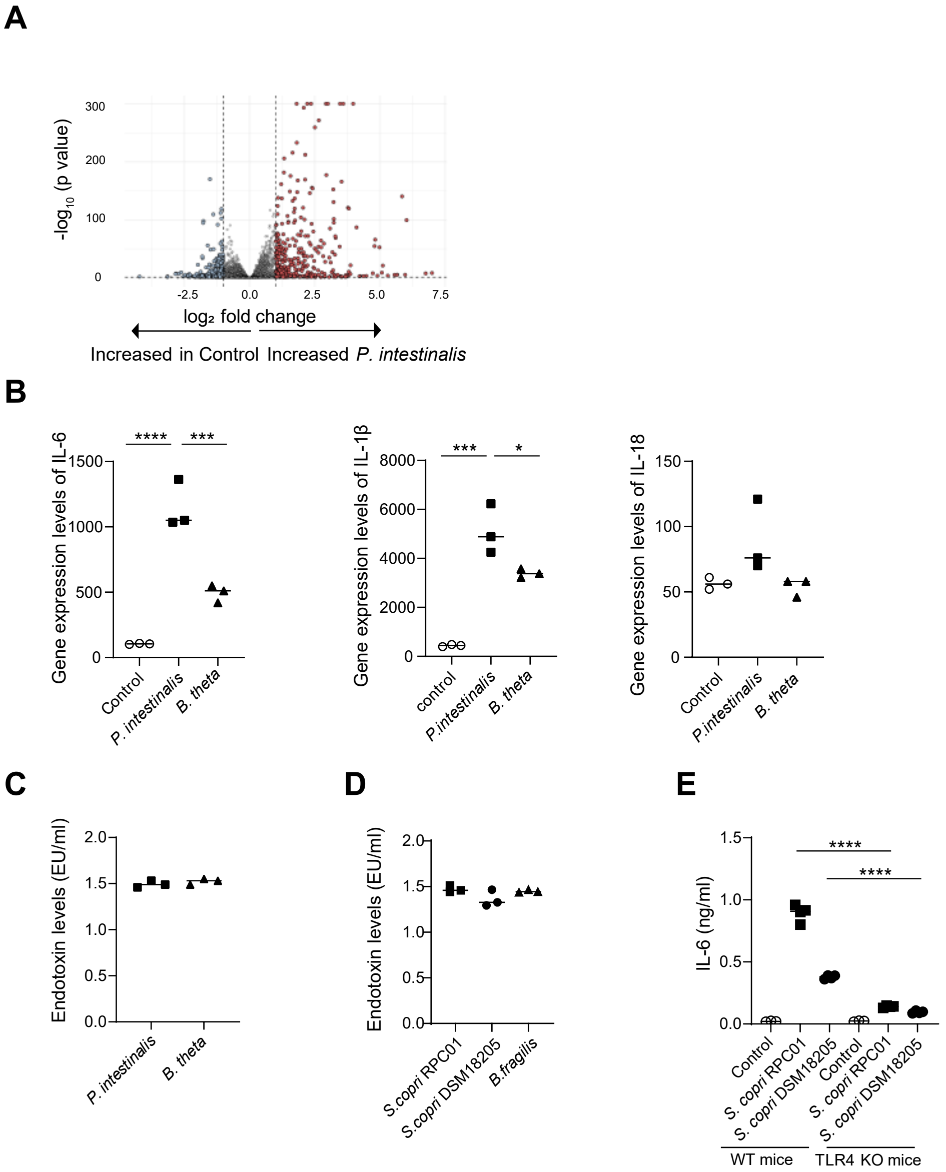


**Figure S8: *P. intestinalis* activates BMDCs and induces IL-6.**
(A) Volcano plot of differentially expressed genes in BMDCs stimulated with *P. intestinalis* vs. control. (B) Gene expression levels of *IL-6*, *IL-1β*, and *IL-18* in BMDCs stimulated with *P. intestinalis*, *B. thetaiotaomicron* (*B. theta*), or control. Upregulated genes in *P. intestinalis*-stimulated DCs are represented by red dots, while downregulated genes are in blue dots. (C) Endotoxin levels in *P. intestinalis* vs. *B. theta*. (D) Endotoxin levels in *S. copri* RPC01, *S. copri* DSM18205, and *B. fragilis*. (E) IL-6 levels in the supernatant of BMDCs from WT and TLR4-KO (TLR4) mice cocultured with *S. copri* RPC01, *S. copri* DSM18205, or control.

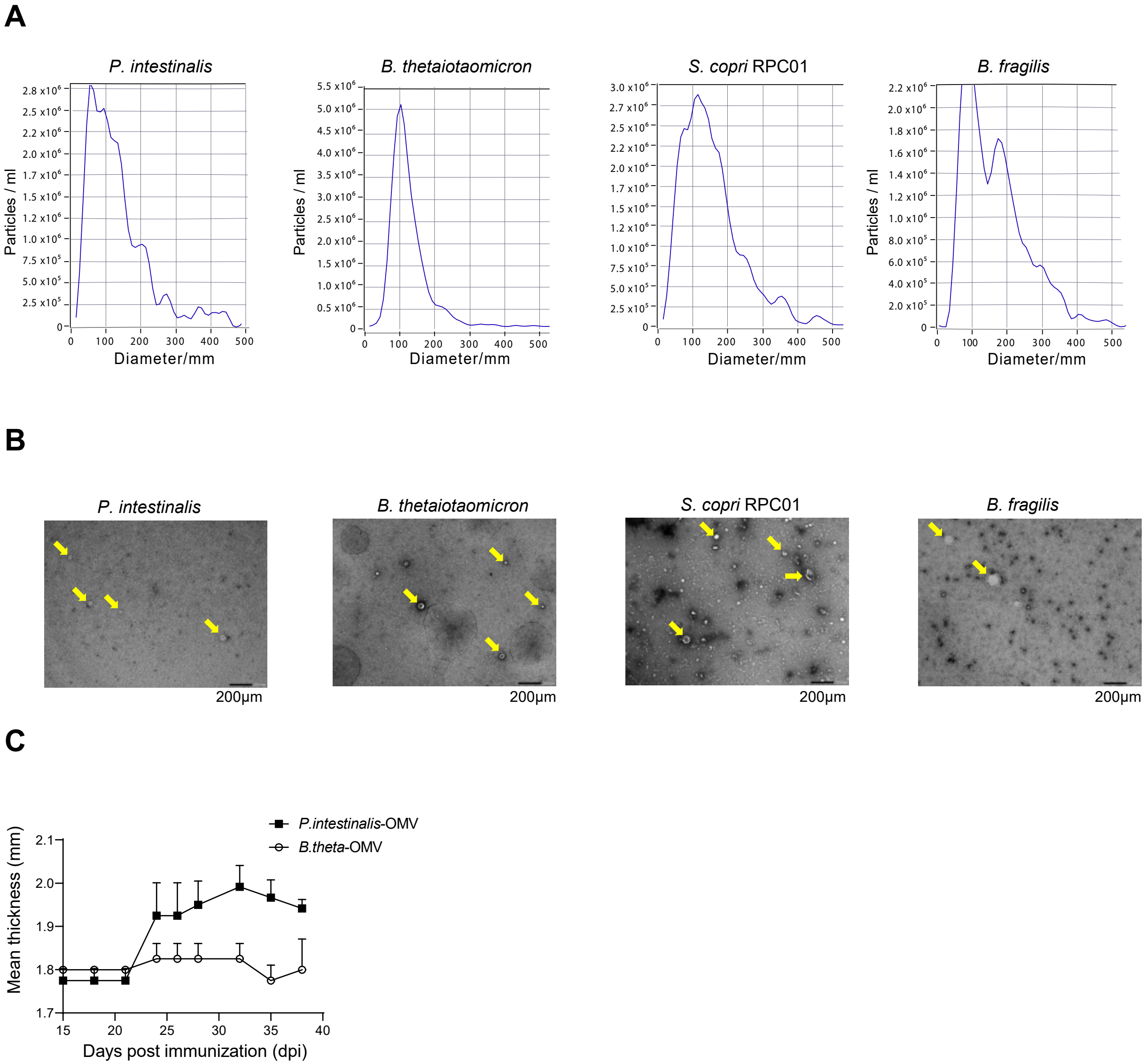


**Figure S9: *Prevotellaceae-derived* OMV induces CIA**

(A) Representative nanoparticle tracking analysis (NTA) of OMVs derived from *P. intestinalis*, *B. theta*, *S. copri* RPC01, and *B. fragilis*. (B) TEM images of OMVs derived from *P. intestinalis*, *B. theta*, *S. copri* RPC01, and *B. fragilis*. Yellow arrows indicate OMVs of the strains. (C) Mean paw thickness in CIA mice inoculated with *P. intestinalis*-OMV vs. *B. theta*-OMV.

**Supplementary Table 1. Clinical characteristics of patients who underwent ileum biopsy**

| Group | New onset RA (n=8) | Established RA (n=10) | Healthy controls (n=9) |
|---|---|---|---|
| Age (years) | 61.22±3.3 | 61.3±2.9 | 62.4±2.9 |
| Female sex (%) | 66.7 | 88.9 | 66.7 |
| Disease duration (years) | 0.4±0.2 | 13.5±3.5 | - |
| RF positivity (%) | 100 | 100 | - |
| ACPA positivity (%) | 88.9 | 100 | 0 |
| DAS28-ESR | 4.95±1.17 | 4.48±1.13 | <0.001 |
| CRP (mg/L) | 2.6±0.7 | 4.0±0.9 | 2.2±0.4 |
| ESR(mm/h) | 11.0±2.4 | 17.3±3.1 | 9.6±4.2 |
| PSL usage (%) | 50.0 | 88.9 | 0 |
| MTX usage (%) | 100 | 80.0 | 0 |
| Rituximab (%) | 0 | 50.0 | 0 |
| Other cDMARD (%) | 11.1 | 40.0 | 0 |

Values represent mean ± standard error (SE), unless otherwise noted. RF = rheumatoid factor, ACPA=anti-cyclic citrullinated peptide antibody, DAS28-ESR = Disease Activity Score in 28 joints using erythrocyte sedimentation rate, PSL = prednisolone, MTX = methotrexate, cDMARD = conventional disease-modifying anti-rheumatic drugs.

**Supplementary Table 2. Relative abundance of *Prevotellaceae* in the human gut**

| Groups | Sample ID | Relative abundance of *Prevotellaceae* (%) |
|---|---|---|
| New onset RA | 14 | 0 |
| | 16 | 0.722 |
| | 20 | 0.083 |
| | 22 | 0 |
| | 24 | 0.068 |
| | 29 | 0 |
| | 42 | 0.075 |
| | 43 | 0 |
| | 49 | 0 |
| Established RA | 1 | 9.493 |
| | 2 | 0.846 |
| | 5 | 0 |
| | 7 | 0 |
| | 8 | 0 |
| | 9 | 0 |
| | 10 | 0 |
| | 11 | 0 |
| | 32 | 0 |
| | 37 | 0.071 |
| Healthy control | 3 | 0 |
| | 6 | 0.072 |
| | 17 | 0.039 |
| | 19 | 0 |
| | 23 | 0 |
| | 25 | 0.029 |
| | 27 | 1.92 |
| | 39 | 0 |
| | 44 | 0.013 |
| | 48 | 0 |

**Supplementary Table 3. Antibodies for flow cytometry**

| Reagent | Source | Identifier |
|---|---|---|
| B220 BV785 Clone RA3-6B2 | Biolegend | Cat# 103246 RRID:AB_11218795 |
| CD11b Alexa Fluor700 Clone M1/70 | Biolegend | Cat# 101222 RRID:AB_493705 |
| CD11c PE -Dazzle594 Clone N418 | Biolegend | Cat# 117348 RRID:AB_2563654 |
| CD19 AF700 Clone 1D3/CD19 | Biolegend | Cat# 152414 RRID:AB_2922474 |
| CD25 SparkNIR685 Clone PC61 | Biolegend | Cat# 102069 RRID:AB_2888823 |
| CD3e APCFire750 Clone 145-2C11 | Biolegend | Cat# 100362 RRID:AB_2629686 |
| CD4 BV750 GK1.5 | Biolegend | Cat# 100467 RRID:AB_2734150 |
| CD44 BV480 Clone IM7 | BD Biosciences | Cat# 566116 RRID:AB_2739518 |
| CD45 FITC Clone 30-F11 | Biolegend | Cat# 103108 RRID:AB_312972 |
| CD62L PE/Cy7 Clone MEL-14 | Biolegend | Cat# 104418 RRID:AB_313102 |
| CD8 BV570 Clone 53-6.7 | Biolegend | Cat# 100740 RRID:AB_10897645 |
| FOXP3 PE Clone FJK-16s | ThermoFisher | Cat# 12-5773 -82 RRID:AB_465936 |
| IFNy APC Clone XMG1.2 | Biolegend | Cat# 505809 RRID:AB_315403 |
| IL-17A BV421 Clone TC11-18H10.1 | Biolegend | Cat# 506926 RRID:AB_10900442 |
| Ly6G e450 Clone 1A8 | ThermoFisher | Cat# 48 -9668 -82 RRID:AB_2637124 |
| RORyt BV421 Clone Q31-378 | BD Biosciences | Cat# 562894 RRID:AB_2687545 |
| T-bet BV605 Clone 4B10 | Biolegend | Cat# 644817 RRID:AB_11219388 |
